# Unraveling fungal endolysosomal network as a potential target for effective disease control

**DOI:** 10.1101/2024.08.02.606287

**Authors:** Xin Chen, Xiaomin Chen, Yunfei Long, Xiang Tian, Zhenyu Fang, Yakubu Saddeeq Abubakar, Huawei Zheng, Zonghua Wang, Wenhui Zheng

**Affiliations:** State Key Laboratory of Agricultural and Forestry Biosecurity, College of Plant Protection, Fujian Agriculture and Forestry University, Fuzhou, 350002, China; Key Laboratory of Bio-pesticide and Chemical Biology, Ministry of Education, College of Plant Protection, Fujian Agriculture and Forestry University, Fuzhou, 350002, China; Fujian University Key Laboratory for Plant Microbe Interaction, College of Life Sciences, Fujian Agriculture and Forestry University, Fuzhou 350002, China; Ministerial and Provincial Joint Innovation Centre for Safety Production of Cross-Strait Crops, College of Geography and Oceanography, Minjiang University, Fuzhou 350108, China; Department of Biochemistry, Faculty of Life Science, Ahmadu Bello University, Zaria 810281, Nigeria

## Abstract

During host-pathogen interactions, fungal pathogens exploit the endolysosomal trafficking network to fine-tune their responses to host and environmental stimuli, thereby facilitating disease progression. However, the molecular mechanisms underlying the fungal-specific functions of the endolysosomal network require further investigation. Here, we systematically characterized the endolysosomal network in *Fusarium graminearum* using the dynamin-like GTPase FgVps1 as an entry point. Functional analysis revealed that FgVps1 is essential for the release of retromer- and sorting nexin-associated vesicles from endosomes, thereby facilitating the trafficking of v-SNARE protein FgSnc1 and promoting fungal development and pathogenicity. Building on this, we further discovered that the retromer core subunit FgVps35 interacts with sorting nexin FgSnx4 and identified the corresponding interaction interface, which involves residues FgVps35^N383^ and FgSnx4^E373^. In addition, the ESCRT-II component FgVps36 bridges ESCRT-I and -III and interacts with both FgVps35 and FgSnx4, thereby preventing their mislocalization to the vacuole and maintaining endolysosomal trafficking. Notably, we demonstrated that inhibition of FgVps1 function, either by blocking its GTPase activity or by disrupting actin polymerization, effectively impaired endosomal trafficking and attenuates fungal pathogenicity. Altogether, our results uncover key mechanisms underlying the function of fungal endolysosomal network and providing a promising broad-spectrum strategy for controlling phytopathogenic fungi.

## Introduction

Each year, crop losses due to emerging fungal pathogens pose a serious risk to global food security ^1, 2^. Successful colonization of fungal pathogens requires not only the ability to evade or suppress host immune responses, but also tightly regulated coordination of intracellular processes that maintain cellular homeostasis and facilitate rapid adaptation to host physiological signals ^3, 4^. Therefore, elucidating the cellular and molecular strategies that pathogens employ to coordinate homeostasis during host colonization is critical to understanding disease progression and developing effective control strategies.

A key role in maintaining cellular homeostasis under biotic and abiotic stress is played by the endolysosomal network, a highly dynamic transport system in eukaryotes that is spatially and temporally regulated ^5^. Following endocytosis, cargo proteins are transported from the cell surface to the early endosomes, where the sorting process takes place. In this phase, cargos are either recycled or degraded ^5, 6^. Recycling pathways include direct return of cargos to the plasma membrane or transit through the endocytic recycling compartment. In addition, some selected cargos can be retrieved into the trans-Golgi network to subsequently reach the cell surface via the secretory pathway. In contrast, cargos destined for degradation are sorted into intraluminal vesicles within the multivesicular bodies at the late endosomes, which subsequently fuse with vacuoles (or lysosomes in mammalian systems) to facilitate degradation ^6, 7, 8^. This process is controlled by an evolutionarily conserved molecular machinery that includes Rab GTPases, retromer, sorting nexins and the ESCRT (Endosomal Sorting Complex Required for Transport) complex. Despite the conservation of these components across species, there are significant differences in gene copy number, functional specialization and regulatory complexity, reflecting species-specific requirements for endosomal transport and homeostatic control.

There is growing evidence for the importance of the endolysosomal network in the pathogenicity of plant-infecting fungi ^9, 10^. Dysfunction of the endolysosomal network affects fungal development and the pathogenesis of various fungal pathogens ^9, 10^. During host-pathogen interactions, components of the endolysosomal system modulate autophagy, a key process that enables fungal cells to adapt to metabolic changes and environmental stress ^11, 12, 13, 14, 15^. Additionally, fungal pathogens utilize the endolysosomal network to facilitate the secretion of effector proteins during host infection to suppress the host immune response ^16, 17^. Furthermore, the endolysosomal network is closely associated with the production of mycotoxin, the toxic secondary metabolites that are toxic to plants and animals but contribute to the pathogen’s ability to invade host tissues ^18, 19, 20, 21, 22, 23, 24, 25^. Despite its crucial role, the molecular complexity of the endolysosomal network in fungal pathogens remains unclearly understood. In particular, the interplay between core trafficking components and other regulatory proteins within this system is yet to be fully elucidated.

In this study, we investigate the endolysosomal network in *Fusarium graminearum*, the causal agent of Fusarium head blight (FHB), to elucidate the molecular basis of endosomal trafficking during the pathogenesis of the fungus. By immunoprecipitation-mass spectrometry (IP-MS), we identified the dynamin-like GTPase FgVps1 as a common interactor of the sorting nexin FgSnx4 and the retromer core subunit FgVps35. Functional analyses revealed that FgVps1 is essential for endosomal sorting, with the v-SNARE protein FgSnc1 identified as one of its cargo proteins. By using FgVps1 as a regulatory nexus, we systematically analyzed the interactions between the core components of the endolysosomal network and elucidate the mechanisms controlling FgSnc1 transport. In addition to the conserved role of FgVps1 in vesicle release, we identified a fungal-specific interaction between FgSnx4 and FgVps35 and uncovered the crucial role of the ESCRT machinery in maintaining endosomal sorting homeostasis. Moreover, we found that phenothiazines selectively inhibit the GTPase activity of FgVps1 by binding the D239-I247 region of the protein, thereby impairing the fungal growth and pathogenicity. In addition to perturbing the GTPase activity of FgVps1, phenothiazine treatment also disrupts actin polymerization, leading to defects in endocytosis and vesicle trafficking. Collectively, these results provide insight into the complex interplay of the endolysosomal network in phytopathogenic fungi and lay a theoretical foundation for the development of environmentally friendly disease management strategies.

## Results

### FgVps1 is important for the development and pathogenicity of *F. graminearum*

We have previously pointed out the important role of the endolysosomal network in the development and pathogenicity of *F. graminearum*, the causal agent of wheat head blight (Fig. S1A) ^18, 19, 20, 22, 23, 24, 26, 27^. A highly enriched protein, FgVps1 (FGSG_07172), was identified by screening the IP-MS data of the retromer core subunit FgVps35 and the sorting nexin FgSnx4. FgVps1 is a dynamin-like GTPase that belongs to the dynamin superfamily. Structurally, FgVps1 contains three domains: a GTPase (G) domain, a dynamin central region (DM) domain and a dynamin GTPase effector (GED) domain (Fig. S1B). We also performed a phylogenetic analysis of the dynamin superfamily proteins and found that FgVps1 is the only homolog of Vps1 involved in endocytic vesicle scission in fungi (Fig. S1C). To further explore the biological functions of FgVps1, we generated Δ*Fgvps1* mutants and confirmed the gene deletion by Southern blot assay (Fig. S2A). We also generated a complemented strain Δ*Fgvps1*-com by expressing a vector harboring GFP-*FgVPS1* into the Δ*Fgvps1* mutant (Fig. S2B). Phenotypic analyses revealed that the absence of FgVps1 leads to severe defects in the vegetative growth, conidiation and sexual reproduction of the fungus (Fig. S2C to S2F). In addition, compared to the wild type and the complemented strain, the virulence of the Δ*Fgvps1* mutant was significantly reduced (Fig. 1A). Overall, these results indicate that FgVps1 loss-of-function perturbs *F. graminearum* development and pathogenicity.

**Fig. 1.**
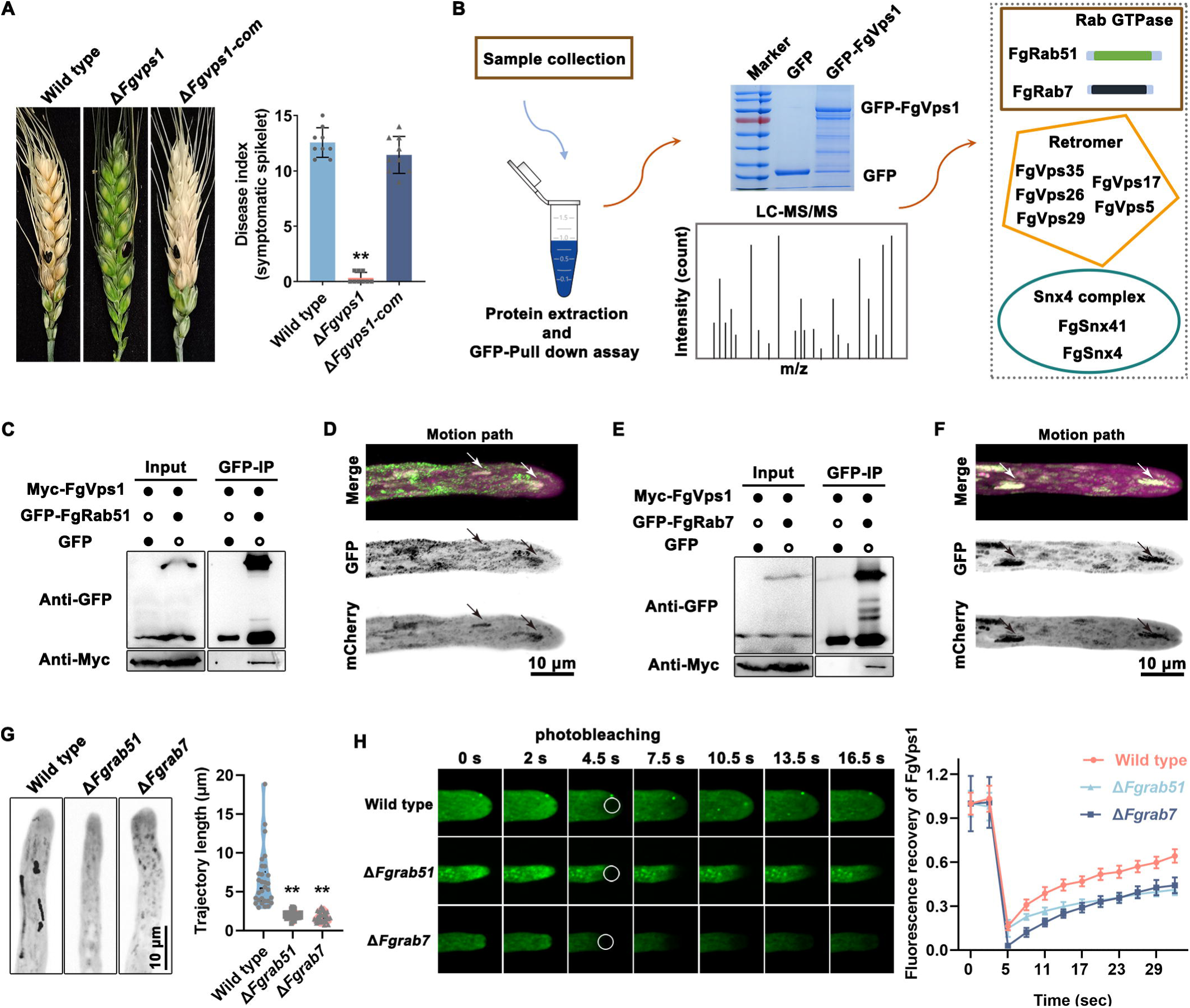
Rab GTPases FgRab51 and FgRab7 coordinate the intracellular dynamic of FgVps1. A. Pathogenicity assay of the indicated strains on wheat heads. B. GFP-FgVps1 pull down strategy for identification of putative FgVps1 interacting proteins. C. Co-IP assay verifying the interaction of FgVps1 with the Rab GTPase FgRab51. D. Time-lapse imaging and motion trajectory analysis of GFP-FgRab51 and mCherry-FgVps1. A total of 37 time-lapse images was taken at 1.64s interval. Arrows indicate shared trafficking trajectory of FgVps1 and FgRab51. E. Co-IP assay verifying the interaction of FgVps1 with the Rab GTPase FgRab7. F. Time-lapse imaging and motion trajectory analysis of GFP-FgRab7 and mCherry-FgVps1. A total of 37 time-lapse images was taken at 1.29s interval. Arrows indicate shared trafficking trajectory of FgVps1 and FgRab7. G. Statistical analysis of the motion trajectories of FgVps1 in the wild type and Rab GTPase mutants (\*\**P*<0.01). The time-lapse imaging involved a total of 150 images was taken at 200 ms interval. H. Fluorescence recovery after photobleaching assay to analyze the fluorescence recovery rate of FgVps1 in the wild type and Rab GTPase mutants.

### The Rab GTPases FgRab51 and FgRab7 regulate the intracellular dynamic of FgVps1

To further decipher the mechanism of FgVps1 in relation to the development and pathogenicity of the fungus, we performed GFP pull-down and subsequent tandem LC-MS/MS mass spectrometry to identify potential FgVps1-associated proteins in *F. graminearum*. As shown in Fig. 1B and Table S3, several proteins that are involved in endolysosomal sorting were detected and identified, including Rab GTPases, the retromer complex and sorting nexins. Along the endolysosomal network, Rab GTPases play a key role in cargo sorting and have been shown to be important for the development and pathogenicity of *F. graminearum* ^6, 27^. Using a Co-IP assay, we found that FgVps1 interacts with FgRab51 (Fig. 1C). To further confirm this association, we co-expressed mCherry-FgVps1 with GFP-FgRab51 in the wild type. Through time-lapse imaging and further trajectory analysis, we found that FgVps1 co-localized with the early endosomal marker FgRab51 (Fig. 1D), consistent with the positive interaction earlier observed. Similar to FgRab51, Co-IP assay and confocal microscopy showed that FgVps1 interacts with FgRab7 and co-localizes with FgRab7-marked late endosomes (Fig. 1E and 1F).

To further investigate the association of FgVps1 with FgRab51 and FgRab7, we transformed a GFP-FgVps1 vector into the Δ*Fgrab51* and Δ*Fgrab7* mutants. Time-lapse analysis revealed that the trajectory length of FgVps1 was significantly reduced in both Δ*FgRab51* and Δ*FgRab7* mutants, suggesting that the Rab GTPases FgRab51 and FgRab7 are required for the proper sub-cellular dynamics of FgVps1 (Fig. 1G). To confirm this result, we conducted a fluorescence recovery after photobleaching (FRAP) assay to monitor FgVps1 diffusion in the respective strains. As shown in Fig. 1H, FgVps1 signal can be recovered by up to 60% within 30s after photobleaching in the wild type, while in the Δ*Fgrab51* and Δ*Fgrab7* mutants, only 40% fluorescence recovery rate was observed. These results demonstrate that the intracellular dynamics of FgVps1 are impaired in the absence of FgRab51 and FgRab7.

In yeast, Mvp1 was shown to recruit Vps1 to the endosomes ^28, 29, 30^. However, in *F. graminearum*, Δ*Fgmvp1* mutant displayed some phenotypic differences with Δ*Fgvps1* ^29, 31^. As shown above, the Rab GTPases contribute to the proper dynamics of FgVps1. Previous studies also demonstrated that Rab GTPases are required for the recruitment of the retromer complex (a protein sorting machinery) and sorting nexins to the endosomes ^16, 19, 26, 32^. Therefore, we proposed that FgRab51 and FgRab7 mediate the recruitment of FgVps1 to the endosomes in *F. graminearum*. To test this hypothesis, we stained the strains expressing GFP-FgVps1 fusion protein with FM4-64 dye to visualize the endosomal structures. Time-lapse imaging revealed that FgVps1 is closely associated with FM4-64-labeled endosomes in the wild-type strain, contrary to their segregated distribution in the Δ*Fgrab51* and Δ*Fgrab7* mutants (Fig. S3). Together with the FRAP assay data, these results further support the important role of Rab GTPases FgRab51 and FgRab7 in maintaining FgVps1 dynamic during endosomal maturation.

### FgVps1 is required for endosomal sorting of FgSnc1

Endocytic cargos travel via the endolysosomal pathway to endosomes where their fate is determined ^5^. Although we have established that FgVps1 is localized to endosomes, its function at the endosomes remains unclear in *F. graminearum*. When screening the FgVps1 IP-MS data, we found that both FgSnx4 and its cargo FgSnc1 were present in the FgVps1 interactome (Fig. 2A, Table S3). Similar to the Δ*Fgsnx4* and Δ*Fgsnc1* mutants, the loss of FgVps1 showed a defect in hyphal morphogenesis (Fig. 2B). Further time-lapse imaging revealed that FgVps1 is closely associated with FgSnc1 in growing hyphae (Fig. 2C). To further confirm the role of FgVps1 in the sorting of FgSnc1, a FRAP assay was performed to investigate the possible diffusion of FgSnc1 in the two mutants Δ*Fgsnx4* and Δ*Fgvps1*. As shown in Fig. 2D, 50% of the FgSnc1 signal in the wild type can be recovered within 1 min after photobleaching, while only 20% fluorescence recovery was observed in the Δ*Fgvps1* and Δ*Fgsnx4* mutants. Furthermore, we also analyzed the localization of FgSnc1 in basal hyphae. Unlike its localization to the plasma membrane in the wild type, FgSnc1 was observed to be mis-localized to the vacuole in both Δ*Fgsnx4* and Δ*Fgvps1* mutants (Fig. 2E). Consistently, western blot assay also showed that the GFP-FgSnc1 band degradation rate was significantly increased in Δ*Fgvps1* and Δ*Fgsnx4* mutants (Fig. 2F). Overall, these results indicate that FgVps1 is required for proper endosomal sorting of FgSnc1.

**Fig. 2.**
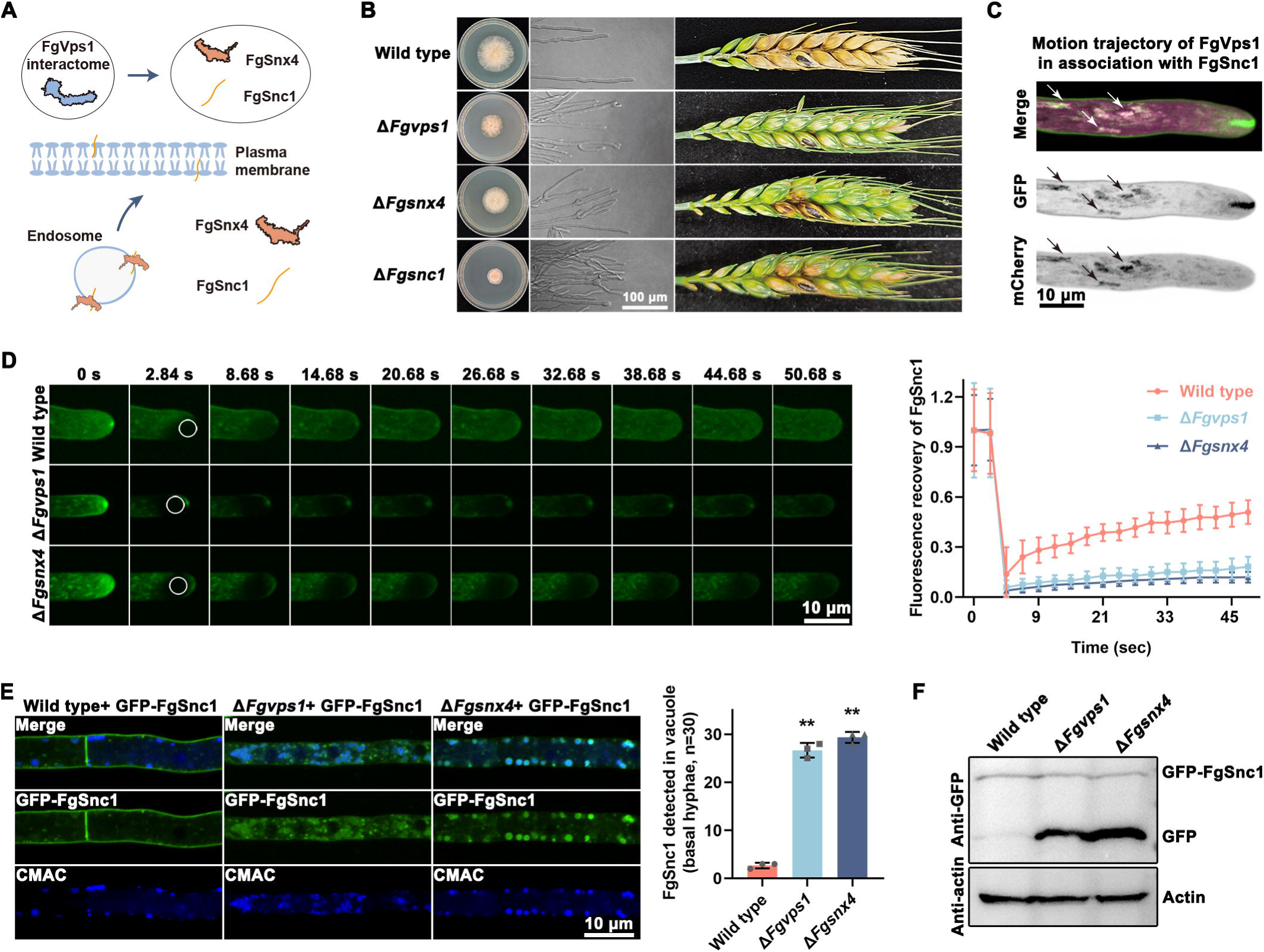
Loss of *FgVPS1* compromises the proper sorting of FgSnc1. A. Prosed model for FgSnx4-mediated FgSnc1 trafficking in *F. graminearum.* B. Analysis of the vegetative growth, apical hyphal morphology and pathogenicity in the indicated strains. C. Time-lapse imaging of GFP-FgSnc1 and mCherry-FgVps1 and analysis of their respective trajectory. A total of 42 images was taken at 1.4s interval. D. Fluorescence recovery after photobleaching assay for the fluorescence recovery rate of FgSnc1 in the wild type, Δ*Fgsnx4* and Δ*Fgvps1* mutants. E. Microscopic observation and statistical analysis of FgSnc1 localization in the basal hyphae of the wild type, Δ*Fgsnx4* and Δ*Fgvps1* mutants. To visualize the vacuolar lumen, the hyphae were stained with CMAC dye. Localization differences were analyzed from 30 hyphae per biological replicate, with three independent replicates involved (\*\**P*<0.01). F. Western blots for FgSnc1 degradation in the wild type, Δ*Fgsnx4* and Δ*Fgvps1* mutants. For western blot analysis, the indicated fungal strains were grown in CM liquid medium for 36 hours.

### FgVps1 is indispensable for the endosomal discharge of FgVps35 and FgSnx4

In *F. graminearum,* both FgVps35 and FgSnx4 contribute to endosomal sorting of FgSnc1 ^19, 33^. To further investigate the role of FgVps1 in the sorting of FgSnc1, Co-IP assay was performed to check for possible interaction between FgVps1 and FgVps35, as well as between FgVps1 and FgSnx4. As shown in Fig. S4A, FgVps1 can interact with FgVps35 and FgSnx4 *in vivo*. Similar to FgVps35 and FgSnx4, FgVps1 can also interact with FgSnc1 (Fig. S4B). Consistent with this, confocal examination also showed that the dynamics of FgVps1 are closely associated with FgVps35 and FgSnx4 (Fig. 3A). Further analysis of the IP-MS data of FgVps1, FgVps35 and FgSnx4 revealed that these proteins share many potential interaction partners (Fig. 3B). Previously, Vps1 was reported to mediate the scission of endosomal tubules in yeast ^30^. To test whether this holds true in *F. graminearum*, we transformed FgSnx4-GFP and FgVps35-GFP vectors into the wild type and the Δ*Fgvps1* mutant. We found that, unlike the wild type, FgVps35 and FgSnx4 were trapped in some speckled structure in the tips and in elongated tubules of the fungal basal hyphae, suggesting the conserved roles of FgVps1 in the endosomal discharge of retromer (FgVps35) and the sorting nexin FgSnx4 (Fig. 3C).

**Fig. 3.**
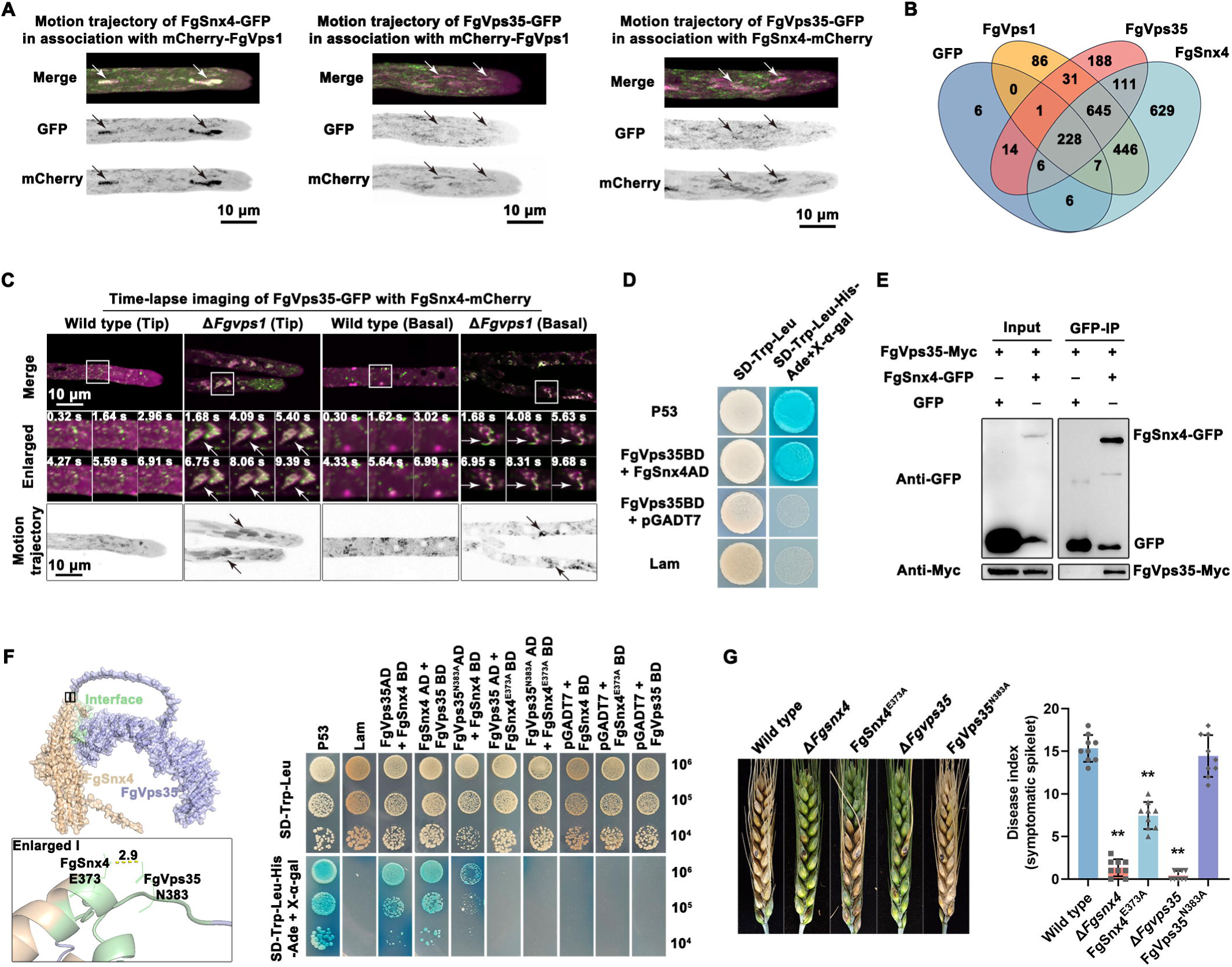
FgVps1 modulates the endosomal release of FgVps35 and FgSnx4. A. Motion trajectory analysis of FgVps1 with FgSnx4 and FgVps35, as well as FgSnx4 with FgVps35. A total of 36 images was taken at 1.6s interval. B. Venn diagram visualization of the interacting proteins of FgVps1, FgVps35 and FgSnx4. C. Time-lapse imaging and trajectory analysis of FgVps35 and FgSnx4 in the wild type and Δ*Fgvps1* mutant. A total of 45 images was taken at 1.3s interval. Arrows indicate speckled structures at the hyphal tips or elongated tubules in the basal hyphae as observed in the Δ*Fgvps1* mutant. D. Yeast two-hybrid assay confirming the interaction between FgVps35 and FgSnx4. E. Co-IP assay verifying the interaction between FgVps35 and FgSnx4. F. Yeast two-hybrid assay verifying the interaction sites of FgVps35 and FgSnx4. The interaction model of FgVps35 with the sorting nexin FgSnx4 was predicted by Alphafold2. FgVps35 is represented in light blue, FgSnx4 is represented in pale green. Black box indicates the zoomed area. Yellow dash represents the hydrogen bond formed between FgSnx4 E373 and FgVps35 N383. G. Analysis of the pathogenicity of the indicated strains on wheat spikes (\*\**P*<0.01).

To exert its function, retromer interacts with various sorting nexins, including Snx1/2, Snx3, Snx5/6 and Snx27 ^34, 35^. However, *F. graminearum* does not have a homolog of Snx27. Moreover, Δ*Fgsnx3* mutant showed phenotypes that are different from those of retromer mutant ^31^. Given that FgSnx4 had similar interactome to FgVps35, we hypothesized that FgVps35 and FgSnx4 may interact to jointly execute their biological functions. To test this hypothesis, yeast two-hybrid and Co-IP assays were carried out to check for possible interaction between FgVps35 and FgSnx4. As shown in Fig. 3D and 3E, FgVps35 directly interacts with FgSnx4 both *in vivo* and *in vitro*. To further validate the association of FgVps35 and FgSnx4, an *in silico* interaction model of FgVps35 and FgSnx4 was analyzed using AlphaFold2. As shown in Fig. 3F, the asparagine residue N383 of FgVps35 was observed to interact with the glutamate residue E373 of FgSnx4 via hydrogen bonding. Sequence alignment further revealed that the E373 of FgSnx4 is evolutionarily conserved across species, and that both E and threonine (T) residues at this position are capable of forming hydrogen bonds, whereas the N383 site appears to be fungi-specific (Fig. S5). To validate this prediction, we disrupted this hydrogen bond by introducing N383A and E373A point mutations in FgVps35 and FgSnx4, respectively, and then analyzed the mutated proteins for possible interaction by yeast two-hybrid (Y2H) assay. We found a weakened interaction between FgVps35 and FgSnx4 due to N383A mutation, and the interaction was completely abolished following E373A mutation. Moreover, pathogenicity tests on wheat heads revealed that the FgSnx4 E373A mutation caused reduced virulence compared to the wild type (Fig. 3G). These results suggest that the FgSnx4 E373 residue is important for the interaction of FgSnx4 with FgVps35.

### The ESCRT complex ensures the continuous sorting of FgSnc1

Previous studies indicated that ubiquitination of Snc1 is crucial for its recycling ^36, 37^. However, although the ubiquitination site of Snc1 remains conserved in *F. graminearum* (Fig. S6 A), mutation of K65 residue of FgSnc1 has no significant effect on the recycling of the protein and does not affect the growth of the fungus, indicating that the FgSnc1 K65 site is not necessary for the sorting of FgSnc1 (Fig. S6 B to E). Within the endosomal network, the ESCRT complex plays a key role in protein degradation and vesicle budding ^7, 38^. Similar to the retromer and sorting nexins, the ESCRT machinery is also required for fungal development and pathogenicity in *F. graminearum* ^22, 23, 24^. Given that ubiquitination of FgSnc1 at the conserved K65 site appears to be dispensable for its recycling and fungal growth, we speculated that FgSnc1 sorting might instead be regulated by alternative mechanisms. Therefore, we hypothesized that the ESCRT complex could function as a potential regulator of FgSnc1 sorting by interacting with its associated sorting machinery. We first investigated the interaction of ESCRT complex with the retromer core component FgVps35 and the sorting nexin FgSnx4. We found that FgVps35 and FgSnx4 positively interact with several components of the ESCRT complex (Fig. 4A). Among them, the ESCRT-II component FgVps36 physically interacts with both FgVps35 and FgSnx4 and serves as a bridge linking ESCRT-I and ESCRT-III (Fig. 4A and 4B). Through CMAC staining (for visualization of vacuole lumen), we found that loss of *FgVPS36* disrupts the endolysosome homeostasis, with more vacuoles observed in both hyphal tips and basal hyphae (Fig. 4C). In addition, compared to the wild type, more band degradation of the endolysosomal components (including FgVps35, FgSnx4, FgVps1 and FgSnc1) was observed in Δ*Fgvps36* mutant (Fig. 4D). Consistent with this observation, live cell imaging showed that FgVps35 and FgSnx4 are mis-localized to the vacuole in Δ*Fgvps36* mutant (Fig. 4E and 4F). We also expressed GFP-FgSnc1 in the Δ*Fgvps36* mutant to analyze its subcellular distribution. Unlike its plasma membrane localization in the wild type, FgSnc1 was highly accumulated in the vacuole in the Δ*Fgvps36* mutant (Fig. 4G to 4I). Altogether, our results indicate that the ESCRT component FgVps36 interacts with the retromer and sorting nexin to modulate the cellular homeostasis of FgSnc1.

**Fig. 4.**
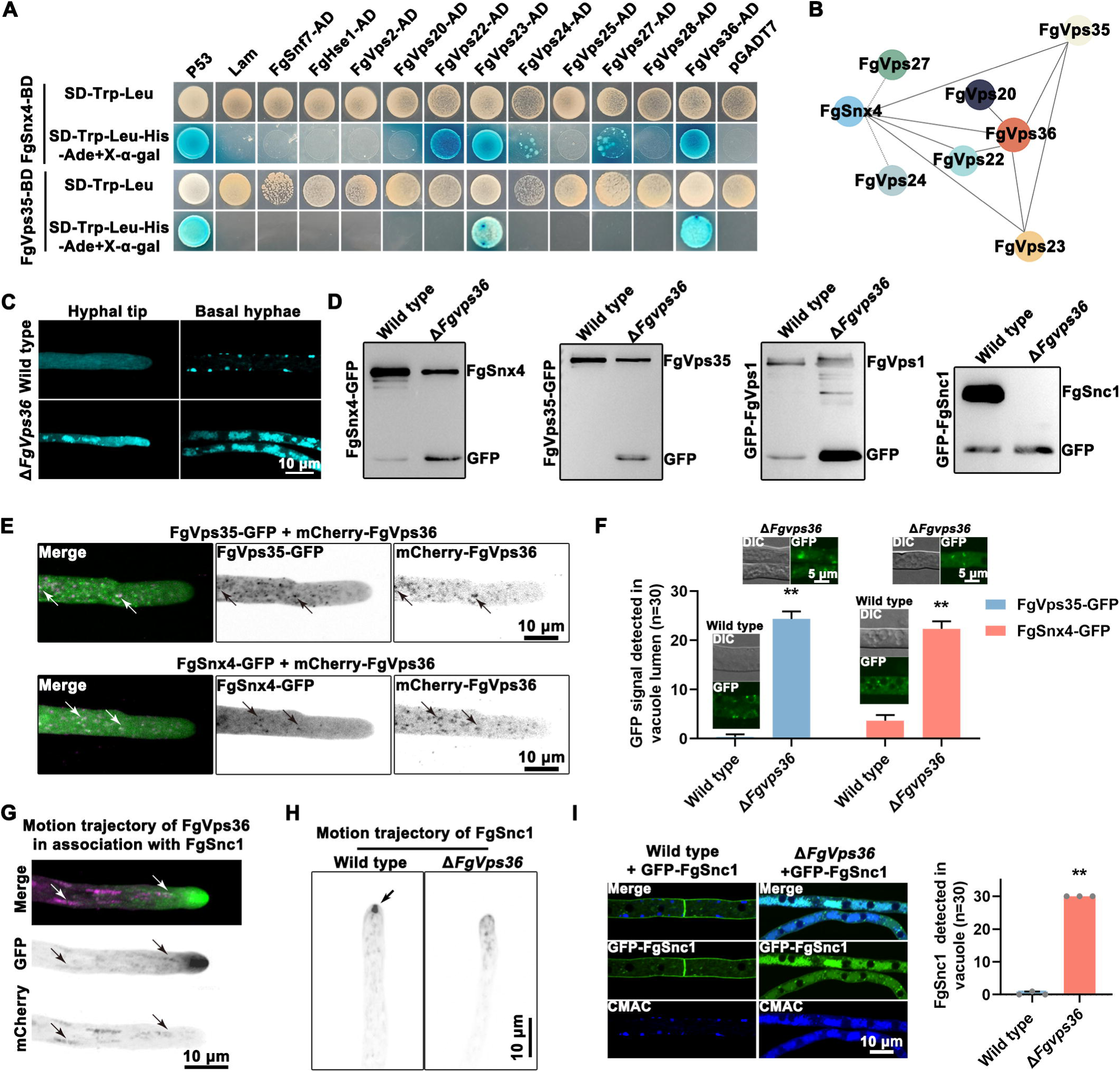
The ESCRT complex component FgVps36 contributes to the homeostasis of the endolysosomal network. A. Yeast two hybrid assay showing positive interaction between the ESCRT machinery and FgVps35 and FgSnx4. B. A simplified schematic showing the interaction among FgVps35, FgSnx4 and the related ESCRT complex components. C. CMAC staining for the observation of vacuole in the wild type and Δ*Fgvps36* mutant. D. Western blot assay for the detection of endosomal network components in the wild type and Δ*Fgvps36* mutant. For western blot assays, the indicated strains were cultured in liquid complete medium for 36 h for proteins extraction, and further enriched using GFP-trap beads. E and F. Effect of *FgVPS36* deletion on the intracellular homeostasis of FgVps35 and FgSnx4. For confocal imaging, the strains were cultured on complete medium for 2 days and then observed under a confocal microscope. Arrows indicate the co-localization of FgVps36 with FgVps35 or FgSnx4. Bar charts represent the quantitative data of the localization patters of FgVps35 and FgSnx4 in the wild type and Δ*Fgvps36* mutant. \*\**P*<0.01. G. Analysis of motion trajectory of FgVps36 with FgSnc1. A total of 12 frames was taken at 5.2-second intervals. Arrows indicate the shared trafficking trajectory of FgVps36 and FgSnc1. H. Analysis of motion trajectory of FgSnc1 in the wild type and Δ*Fgvps36* mutant. A total of 57 frames was taken at 500ms intervals. Black arrow indicates the movement of FgSnc1 at the hyphal tip. I. Confocal examination for the distribution of FgSnc1 in the basal hyphae of the wild type and Δ*Fgvps36* mutant. Vacuole were stained with CMAC dye, \*\**P*<0.01.

### Phenothiazine treatment inhibits FgVps1 GTPase activity and subsequent FgSnc1 sorting

In contrast to other species, Vps1 is the only protein of the dynamin superfamily that is used by phytopathogenic fungi for endocytic transport^39^. To determine whether inhibition of FgVps1 could serve as a potential disease control strategy, the phenothiazine derivatives chlorpromazine (CPZ) and prochlorperazine (PCZ) were used for further treatment. Phenothiazines, a clinically used antipsychotic, exhibits dynamin inhibitory activity, but its effect on phytopathogens remains largely unexplored ^40, 41^. As shown in Fig. S7A, treatment with phenothiazines can significantly impair the growth of *F. graminearum*. We also investigated the effects of phenothiazines on plant infection. In particular, when treated with CPZ and PCZ, *F. graminearum* showed severe defects in invasive growth and virulence on wheat coleoptiles (Fig. S7B and S7C). Since phenothiazines were previously shown to inhibit dynamin GTPase activity, we further purified the recombinant GST-FgVps1 protein and incubated it with phenothiazines for 30 min. Subsequent GTPase activity assay indicated that the FgVps1 GTPase activity was significantly reduced under phenothiazines treatment, indicating the conserved role of phenothiazines in inhibiting the GTPase activity of dynamins (Fig. 5A). In addition, treatment with phenothiazines compromised the normal intracellular distribution of FgVps1 and disrupted the intracellular trafficking of FgSnc1 (Fig. 5B and 5C).

**Fig. 5.**
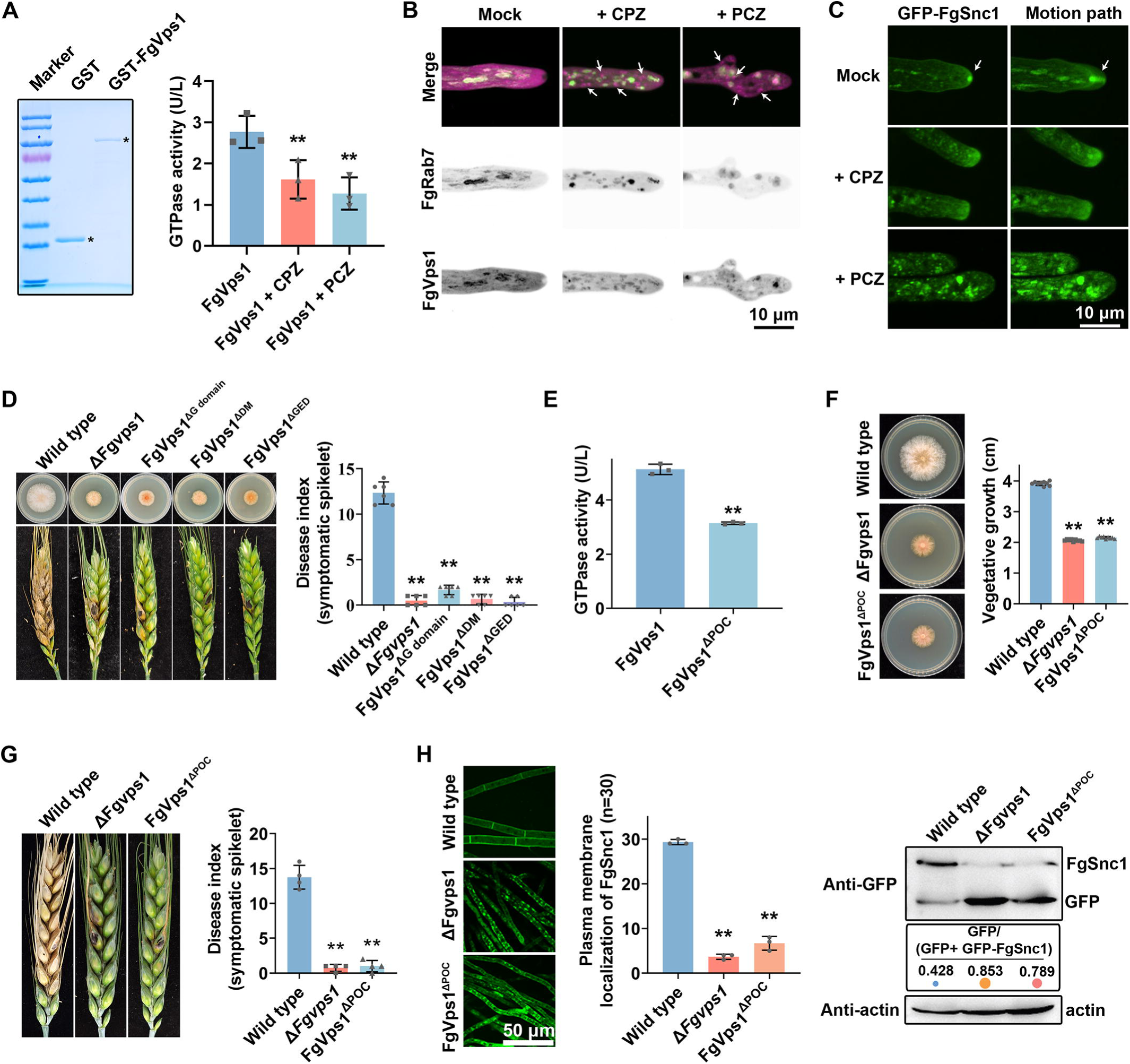
Inhibition of FgVps1 GTPase activity perturbs FgSnc1 sorting in *F. gramienarum*. A. Purification of GST and GST-FgVps1 recombinant proteins and further analysis of GTPase activity. To evaluate the effects of phenothiazines on FgVps1 GTPase activity, 5 μL of the purified GST-FgVps1 was incubated with 5 μL of phenothiazines (at a final concentration of 100 μg/mL) for 30 min and then used for GTPase activity assay. B. Time-lapse imaging and trajectory analysis for the effect of phenothiazines treatment on FgVps1 distribution. For confocal examination, the strains expressing GFP-FgRab7 and mCherry-FgVps1 was inoculated on SYM medium with or without phenothiazines for 2 days. A total of 47 images was taken at 1.2s interval. White arrows indicate the abnormal puncta detected in the cytosol. C. Time-lapse imaging and trajectory analysis for the effect of phenothiazines treatment on FgSnc1 distribution. A total of 300 images was taken at 200ms interval. White arrows indicate the dynamics of FgSnc1 at the hyphal tip. D. Phenotype assays for the effects of FgVps1 domain deletions on fungal development and pathogenicity. E. GTPase activity assays of FgVps1^ΔPOC^. The concentration of the purified recombinant proteins was quantified, and then 10 μL of each protein was used for further GTPase activity assay. F. Growth assay of FgVps1^ΔPOC^ strain on CM medium. G. Pathogenicity assays of FgVps1^ΔPOC^ strain on wheat heads. G. Confocal microscopy and western blot assays for the effect of FgVps1^ΔPOC^ on FgSnc1 sorting (\*\**P*<0.01).

As a typical protein of the dynamin superfamily, FgVps1 contains three domains: G-domain (GTPase domain), DM (dynamin central region) and GED (dynamin GTPase effector domain) (Fig. S1B). By mutating the domains, we found that these domains all contribute to the full function and subcellular distribution of FgVps1 (Fig. 5D and S8A). To further identify the structural basis of FgVps1 association with phenothiazines, we constructed the molecular docking model of FgVps1 with the phenothiazines CPZ and PCZ using AutoDock ^42^. Interestingly, among the different binding pockets identified, CPZ and PCZ share the same binding pocket with FgVps1, which consists of amino acids from D239 to I247 (Fig. S8B). The binding pocket for phenothiazines belongs to the G domain, which is involved in the binding and hydrolysis of GTP ^39^. To determine whether this binding pocket is functionally important for the GTPase activity of FgVps1, we generated a truncated FgVps1 protein lacking this pocket (FgVps1^ΔPOC^, deletion of residues D239-I247). Subsequently, we purified the recombinant GST-FgVps1^ΔPOC^ protein and analyzed its GTPase activity in vitro (Fig. S8C). As shown in Fig. 5E, the mutation of the binding pocket caused a significant reduction in the GTPase activity of FgVps1. Also, loss of the phenothiazines binding pocket impairs the full function of FgVps1 (Fig. 5F and 5G). Consistent with *FgVPS1* deletion and G domain mutation, loss of the phenothiazines binding pocket disrupted the proper FgSnc1 sorting (Fig. 5H). Taken together, these findings suggest that inhibiting the GTPase activity of FgVps1, either by genetic or chemical means, significantly impairs the development and pathogenicity of *F. graminearum*.

### Treatment of phenothiazines impaired actin polymerization in phytopathogenic fungi

It was noted that treatment of *F. graminearum* with a high concentration of phenothiazines (100 μg/mL) caused more severe defects than those observed in the *FgVPS1* deletion mutant, suggesting that apart from disruption of FgVps1 activity, phenothiazines affect additional cellular processes. Considering the central role of actin in promoting cell polarity and vesicle trafficking ^43^, we speculated that phenothiazines could interfere with actin organization. To check this, we treated a strain expressing GFP-FgSnc1 with phenothiazines or the actin polymerization inhibitor Latrunculin A (Lat A). As shown in Fig. S9A, both treatments disrupted the polarized localization of FgSnc1. We further assessed the effect of phenothiazines on actin organization and found that actin localization at the Spitzenkörper was markedly reduced, indicating disrupted polymerization (Fig. 6A). Since actin organization is also crucial for endocytosis in filamentous fungi, we investigated the effects of phenothiazines on this process using FM4-64 dye. In the control hyphae (treated with DMSO), FM4-64 clearly labeled the vacuolar membrane within 25 min. In contrast, FM4-64 was observed to accumulate in the cytosol in the phenothiazine-treated hyphae, indicating delayed endocytosis (Fig. 6B). To further validate these observations, we monitored the internalization of the uric acid-xanthine permease FgUapC ^27^. During growth in MM-N medium containing 5 μM urea, FgUapC mainly localized to the plasma membrane. Upon switching to an ammonium-containing medium (10 μM NH₄Cl), FgUapC was internalized and delivered to the vacuole in the control cells. However, in phenothiazine-treated hyphae, FgUapC remained localized at the plasma membrane even after 3.5 hours, indicating a delay in endocytic trafficking (Fig. 6C).

**Fig. 6.**
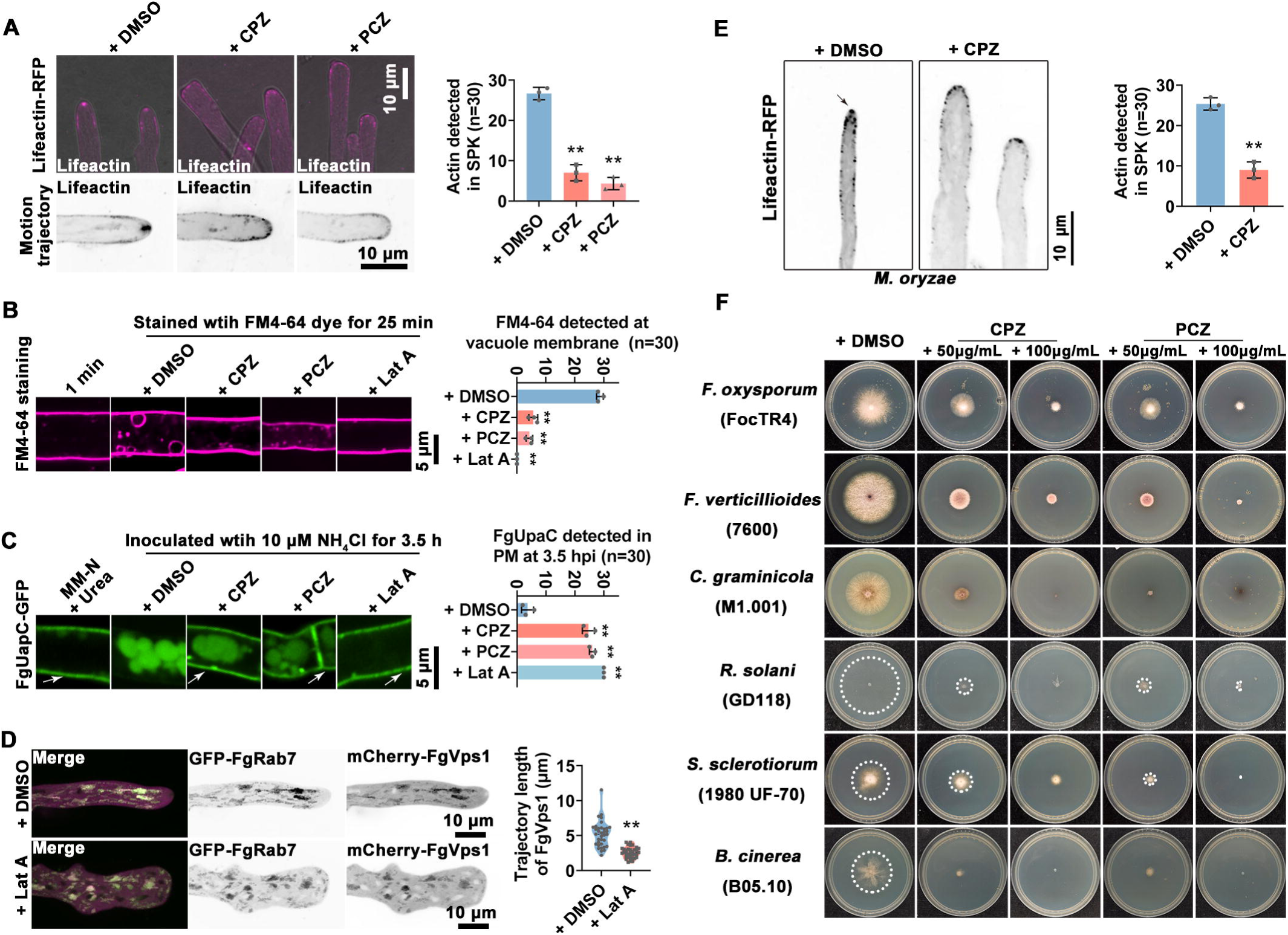
Treatment with phenothiazines disrupts actin organization in SPK. A. Effect of phenothiazine treatment on actin organization at hyphal tip polarity (\*\**P*<0.01). To evaluate the effect of phenothiazines on actin organization, the wild type strain expressing Lifeactin-RFP was inoculated onto CM plates supplemented with or without phenothiazines (50 μg/mL) for 2 days (\*\**P*<0.01). For motion trajectory analysis, A total of 47 images was taken at 400ms interval. B. FM4-64 staining for monitoring endocytosis following treatment with phenothiazines (50 μg/mL) and Lat A (5 μg/mL). C. Confocal examination for the endocytosis of FgUapC under phenothiazine and Lat A treatments. To induce the endocytosis of FgUapC, the wild type strain expressing GFP-FgUapC was shifted from a urea-containing medium to an ammonium-rich medium (10 μM NH₄Cl) for 3.5 hours. Phenothiazines and Lat A were used at final concentrations of 50 μg/mL and 5 μg/mL, respectively. White arrows indicate the plasma membrane localization of FgUapC. D. Confocal examination and motion trajectory analysis of the effects of Lat A treatment on FgVps1 dynamic. A total of 29 images was taken at 1s interval. E. Confocal examination for the effect of CPZ treatment on actin organization in *M. oryzae* (\*\**P*<0.01). To evaluate the effect of the phenothiazine derivative CPZ on actin organization, the wild type strain (70-15) expressing Lifeactin-RFP was inoculated on CM plates containing 50 μg/mL CPZ. F. Effects of phenothiazines treatment on the growth of the indicated fungal pathogens.

Given the important role of actin in vesicle trafficking in filamentous fungi^43^, we then investigated whether actin organization influences FgVps1 dynamics. In a strain co-expressing GFP-FgRab7 and mCherry-FgVps1, Lat A treatment did not alter the endosomal localization of FgVps1 but significantly reduced its dynamic movement as indicated by shorter tracking trajectories (Fig. 6D). To determine whether these effects are conserved across phytopathogenic fungi, we treated *Magnaporthe oryzae* with the phenothiazine derivatives. Consistent with the findings in *F. graminearum*, treatment of *M. oryzae* with the drugs caused defects in the fungal development and pathogenicity (Fig. S9B). Moreover, treatment with CPZ disrupted actin polymerization in *M. oryzae* (Fig. 6E). We also treated other phytopathogenic fungi, including *Fusarium oxysporum*, *Fusarium verticillioides*, *Colletotrichum graminicola*, *Rhizoctonia solani*, *Sclerotinia sclerotiorum*, and *Botrytis cinerea* with the drugs and observed similar growth inhibition effect across these species. Collectively, these results indicate that phenothiazines exert antifungal activity not only by impairing FgVps1-mediated vesicle trafficking but also by disrupting actin polymerization and subsequent endocytosis, ultimately compromising fungal growth and virulence.

## Discussion

Accumulating evidence has highlighted the crucial role of endolysosomal network in maintaining cellular homeostasis and its close association with the development and pathogenesis of phytopathogenic fungi ^5, 10^. However, the intricate mechanisms regulating this network and its potential for fungal disease control remains limited. In this study, we identified the dynamin-like GTPase FgVps1 as a molecular “scissor” functioning in the endosomal network. FgVps1 is first recruited to endosomes by the Rab GTPases FgRab51 and FgRab7, where it facilitates the release of vesicles containing the retromer and sorting nexins. The efficiency of this process is further regulated by the ESCRT complex interacts directly with FgVps35 and FgSnx4. Importantly, disruption of endosomal sorting, either by inhibiting FgVps1 activity or blocking actin polymerization, significantly compromised the growth and virulence of *F. graminearum* (Fig. 7). These findings unveil new insights into the functions of the endolysosomal system and suggest a potentially promising strategy for phytopathogenic fungal disease control.

**Fig. 7.**
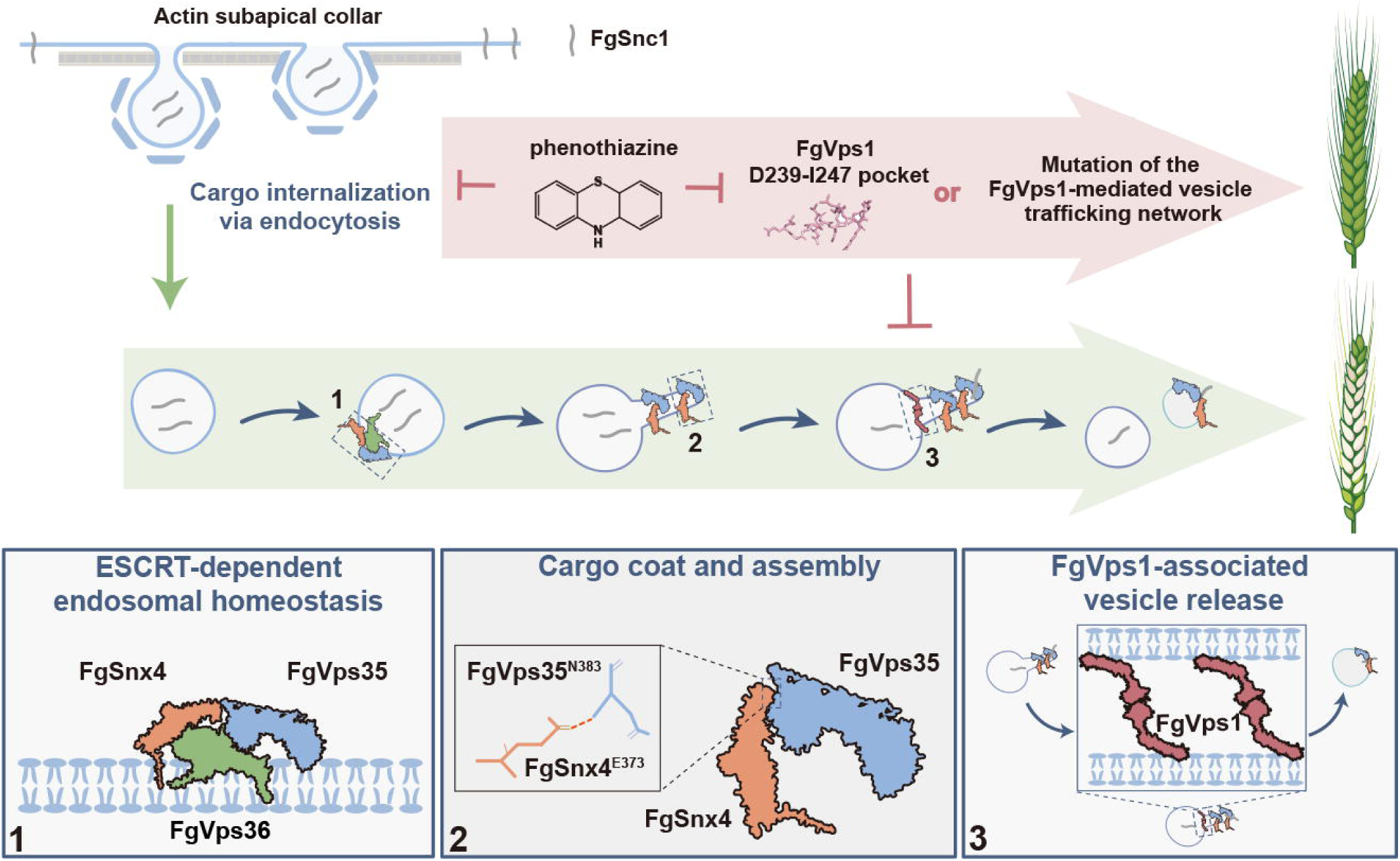
Proposed model for FgVps1-associated endosomal sorting in *F. graminearum*. Through endocytosis, cargo proteins (e.g. FgSnc1) are internalized and delivered to endosomes, where endosomal sorting is initiated. At the endosomes, sorting nexins and the retromer complex recognize specific cargo proteins and facilitate the formation of endosomal tubules. During this process, the ESCRT machinery interacts with the sorting nexins and the retromer to prevent misrouting and premature degradation of the cargos. As endosomal tubules mature, Rab GTPases recruit the dynamin-like GTPase to promote vesicles release. The released vesicles are then trafficked to target membranes for subsequent recycling. Disruption of the endosomal sorting, either by inhibition of FgVps1 function or disruption of actin organization, efficiently inhibit fungal development and pathogenicity.

Dynamins are multi-domain GTPases involved in multiple membrane remodeling processes (including scission and fusion of endocytic vesicles and organelles) and cell division events ^39, 44, 45^. Unlike the many dynamin proteins identified in mammals and plants ^39, 44^, fungi contain only three dynamin proteins namely Dnm1, Vps1 and Sey1; Dnm1-dependent mitochondrial fission and Sey1-mediated ER fusion have been shown to be important for the development, fungicide resistance and pathogenicity of phytopathogens ^46, 47, 48^. Although Vps1 has previously been linked to endocytosis, vesicle scission and peroxisome fission and recycling ^29, 30, 49, 50, 51, 52^, its precise roles in fungal phytopathogens remain poorly understood. In this study, we demonstrated that FgVps1 plays a conserved role in vesicle release and provided compelling evidence that its function and dynamics are dependent on Rab GTPases and proper actin organization (Fig. 1, Fig. 3, and Fig. 6). Previous studies in yeast have shown that the endosomal recruitment of Vps1 depends on Mvp1, which facilitates the release of recycling tubules ^28, 29, 30^. However, in *F. graminearum*, there are clear phenotypic differences between Δ*Fgmvp1* and Δ*Fgvps1* mutants ^31^, suggesting that FgVps1 exerts its functions independent of FgMvp1. Our results demonstrated that the Rab GTPases FgRab51 and FgRab7 contribute to the endosomal recruitment of FgVps1 and are essential for maintaining its intracellular dynamics (Fig. 1 and Fig. S3). FgVps1 function, Rab GTPases also play a conserved role in recruiting sorting nexins and the retromer to endosomes, thereby coordinating the endosomal sorting process ^19, 26, 32, 53^. The functional diversity of Rab GTPases reflects an evolutionary adaptation that ensures precise regulation of endosomal trafficking during pathogenic fungal development and host colonization. Additionally, our findings demonstrate that proper actin organization also contributes to the dynamic of FgVps1 (Fig. 6). Given the crucial role of actin in both endocytosis and vesicle trafficking ^5, 43, 54^, disrupting endolysosomal homeostasis by interfering with fungus-specific actin-driven vesicle motility may offer a promising strategy for fungal disease management.

Moreover, we also systematically characterized the role of FgVps1 in endosomal sorting using FgSnc1 as a representative cargo. Intriguingly, we found that FgSnx4 and FgVps35 are trapped in the same endosomal tubule in Δ*Fgvps1* mutant (Fig. 3C). Further analysis revealed that FgSnx4 directly interacts with FgVps35 through the E737 residue of the former (Fig. 3F). Previous studies in yeast and mammals demonstrated that the retromer interacts with different sorting nexins to modulate cargo sorting ^35, 55^; however, some of those sorting nexins are either absent or their loss-of-function displays different phenotypes in phytopathogens. The observed interaction between FgSnx4 and FgVsps35 may thus represent an evolved adaptation in phytopathogens to optimize intracellular trafficking for successful host colonization. To better understand the specialized endosomal sorting mechanisms in these pathogens, we attempted to investigate the relationship between the FgSnx4-FgVps35 heterodimer and the ESCRT complex. We found that the ESCRT complex is crucial for proper cargo sorting and directly interacts with both FgSnx4 and FgVps35 to maintain endosomal homeostasis (Fig. 4). Interestingly, unlike the compartmentalized network seen in yeast and mammals, the endosomal system in fungal pathogens appears to be structurally “coarse” yet functionally efficient. This functional robustness is likely achieved through the tight interplay of Rab GTPases (Rab5 and Rab7), dynamin (FgVps1), sorting machineries (retromer and sorting nexins) and the ESCRT complex (Fig.1, Fig. 3 and Fig. 4) ^19, 26, 29, 31, 56^. Together, these findings provide new insights into the role of the endolysosomal network in phytopathogens and its contribution to fungal development and host infection.

The antipsychotic drugs phenothiazines exhibit antimicrobial effects against human pathogenic bacterial and fungi ^41, 57, 58^. Previous studies indicated that phenothiazines interfere with calcium-calmodulin system, membrane stabilization and efflux pumps to strive an antifungal effect ^59, 60^. In this study, we provide new evidence that phenothiazines exert antifungal effects by compromizing the function of the endolysosomal network. Specifically, phenothiazines target and bind to D239-I247 amino acid residues in the GTPase (G) domain of FgVps1, thereby inhibiting its GTPase activity and disrupting FgVps1-associated endosomal sorting (Fig. 5). In parallel, phenothiazine treatment interferes with actin polymerization in phytopathogenic fungi, leading to defects in polarity establishment and endocytosis. This disruption also impairs the intracellular dynamics of FgVps1, further compromising endolysosomal trafficking (Fig. 6). To facilitate host colonization, many pathogens manipulate the host’s endolysosomal system to enhance their colonization efficiency ^61, 62, 63, 64^. it is plausible that phenothiazines may also inhibit pathogen entry by targeting host endocytosis. Since phenothiazines are synthetic drugs with diverse biological activities ^57^, further understanding of their mechanism of action may be promising for effective drug development and repurposing.

## Materials and methods

### Fungal strains and culture conditions

The PH-1 strain of *F. graminearum* was used as the wild type. A list of all the strains used in this study is provided in Table S1. The strains were cultured as previously described ^18^. In brief, the *F. graminearum* strains were cultured on CM and CMC media for growth and conidia production, respectively.

To assess the impact of phenothiazines on the indicated fungal pathogens, *Fusarium oxysporum* (FocTR4) and *Fusarium verticillioides* (7600) were inoculated on CM media, while *Colletotrichum graminicola* (M1.001), *Rhizoctonia solani* (GD118), *Sclerotinia sclerotiorum* (1980 UF-70) and *Botrytis cinerea* (B05.10) were inoculated on CMII media (10 g/L glucose, 2 g/L peptone, 1 g/L yeast extract, 1 g/L casamino acid, 5% (v/v) nitrate salts, 0.1% (v/v) trace elements and 0.1% (v/v) vitamin solution). After inoculation, all strains were incubated at 28 °C under dark condition.

### Generation of *FgVPS1* gene deletion mutants and complementation

For the generation of Δ*Fgvps1* mutants, split-marker approach was used as described previously ^65^. Briefly, approximately 1kb upstream and downstream sequences of *FgVPS1* gene were amplified using specific primer pairs (Table S2). The resulting fragments were then fused with hygromycin resistance gene cassette through splicing by overlap extension (SOE) PCR and then transformed into the wild type PH-1 protoplasts. The resulting transformants were screened by PCR and further confirmed by Southern blot assay. For *FgVPS1* complementation, a GFP-Fgvps1 vector was transformed into the Δ*Fgvps1* mutant protoplasts and, subsequently, positive transformants were screened by GFP fluorescence detection using a confocal microscope as described below. Additionally, the selected transformants were further verified through phenotypic assays.

### Plant infection assays

For *F. graminearum* infection on wheat coleoptiles, conidia suspensions (10^6^ spores/mL) of the indicated strains were inoculated on wheat coleoptiles for 7 days as previous descripted ^66^. For pathogenicity on wheat heads, mycelial plugs of the related strains were inoculated into flowering wheat heads for 14 days as previously descripted ^18^. To monitor invasive growth, conidia suspensions were injected into wheat coleoptiles and incubated in the dark for 3 days, under humid condition at 28 ℃, and then observed under a confocal microscope.

For *M. oryzae* pathogenicity assays on barley, conidia suspensions (5 × 10⁴ conidia/mL) were drop-inoculated onto barley leaves with or without phenothiazine treatment and incubated for 5 days, as previously described ^65^.

### Confocal microscopy and analysis

For monitoring endocytosis, hyphae of the fungal strains were incubated with 10 μg/mL FM4-64 dye (Cat. T3166, Invitrogen, Carlsbad, California) for 2 and 25 minutes, respectively. To stain the vacuoles, hyphae of the indicated strains were treated with 50 μM CMAC (Cat. A6520, Invitrogen, Carlsbad, California) for 30 min and then observed under a confocal microscope. For the disruption of action polymerization, Latrunculin A (Lat A) was used at a final concentration of 5 μg/mL and observed by confocal microscopy. All confocal microscopy examinations were performed using a Nikon CSU-W1 spinning disk confocal microscope (Nikon, Japan). The excitation and emission wavelengths used were 488 nm/500-550 nm for GFP and 561 nm/570-620 nm for mCherry as previously descripted ^16^.

For statistical analysis of localization differences, data were collected from 30 hyphae per biological replicate across three independent replicates. For the analysis of vesicle motion, time-lapse image stacks were processed using Fiji ^67^. Maximum intensity projections (MIPs) were generated from the image stack to visualize the vesicle trajectory across the time series. Single-particle tracking analysis was then performed using a TrackMate plugin. The net displacement of individual vesicles was calculated as the straight-line distance between their initial and final positions, based on the trajectories obtained from the single-particle tracking.

### Immunoprecipitation-mass spectrometry

For the immunoprecipitation-mass spectrometry assays, the wild type strain PH-1 and the strains expressing GFP and GFP-FgVps1 were cultured in liquid CM for two days. Mycelia were harvested, ground into fine powder and then incubated in lysis buffer containing 2 mM PMSF and protease inhibitor cocktail (Cat. GRF101, Epizyme Biotech, Shanghai, China) for 30 min. The samples were centrifuged at 12,000 rpm for 20 minutes. Total cell lysates were further incubated with 30 μL GFP-Trap A beads (Cat. gta-20; ChromoTek Inc., Planegg Martinsried, Germany) for 4 hours and then heated at 100℃ with protein loading buffer (Cat. LT103, Epizyme Biotech, Shanghai, China) for 10 minutes. Finally, the samples were separated by SDS-PAGE, and mass spectrometry analysis was conducted to identify the related proteins bound to the GFP-FgVps1.

### Yeast two hybrid and co-immunoprecipitation (Co-IP) assays

For yeast two hybrid assays, the CDS sequence of the various genes were amplified using the specific primers listed in Table S2. The amplified sequences were cloned into pGADT7 or pGBKT7 vector and further verified by sequencing (Sangon biotech, Shanghai, China). The resulting vectors were transformed into yeast strain AH109 according to Matchmaker™ GAL4 Two-Hybrid System 3 protocol (Clontech, Mountain View, CA, USA).

For Co-IP assay, the strains expressing the fusion proteins were inoculated in liquid CM at 28℃, 110 rpm for 36 h. The samples were collected and ground into fine powder in liquid nitrogen, and then lysed in lysis buffer. The total cell lysates were then incubated with GFP-Trap A beads at 4 ℃ for 4 h. The bound proteins were subsequently eluted by heating at 100℃ for 10 min with protein loading buffer and further analyzed by western blot using anti-GFP (Cat. M20004M, Abmart, Shanghai, China) and anti-Myc (Cat. ab1326, Abcam, Cambridge, United Kingdom) antibodies.

### Phylogeny analysis, protein structure modeling and molecular docking

For phylogeny analysis, the sequences of the dynamin proteins were downloaded from the NCBI database (https://www.ncbi.nlm.nih.gov/). Phylogenetic tree was constructed by Neighbor-joining method using MEGA-X and further modified using iTOL ^68^. The protein structures of FgVps1 and FgVps35-FgSnx4 multimer were predicted using AlphaFold2 ^69^ and further visualized using PyMOL. The structures of prochlorperazine and chlorpromazine were downloaded from the NCBI PubChem database and molecular docking of FgVps1 with the prochlorperazine and chlorpromazine was performed using AutoDock and visualized by PyMOL.

### Protein purification and GTPase activity assay

For the purification of GST-FgVps1 and GST-FgVps1^ΔPOC^ recombinant proteins, the CDS sequences of *FgVPS1* and *FgVPS1*^ΔPOC^ were amplified using the specific primers listed in Table S2 and then cloned into pGET-4T-2 vector. The resulting vectors were expressed in *Escherichia coli* BL21 cells and further sonicated in lysis buffer (150 mM NaCl, 50 mM Tris HCL, pH 8.0, 10% glycerol) containing 2 mM PMSF and protease inhibitor cocktail (Cat. GRF101, Epizyme Biotech, Shanghai, China). After sonication, the suspensions were centrifuged at 4 ℃, 12000rpm for 20 min and incubated with GST beads (Cat. A10018, Abmart, Shanghai, China) for 2 h. The beads were then washed with the washing buffer and eluted by incubating in an elution buffer (150 mM NaCl, 50 mM Tris HCL, pH 8.0, 150 mM glutathione) at 4 ℃ for 2 h.

To investigate the effects of phenothiazines on FgVps1 GTPase activity, the purified GST-FgVps1 was incubated with the different phenothiazines for 30 min and then the GTPase activity was measured using a GTPase assay kit (Cat. ab272520, Abcam, Cambridge, United Kingdom) and Tristar² S LB 942 multimode reader (Berthold, Germany). To check the effect of mutating the binding pocket of FgVps1 on its GTPase activity, the purified GST-FgVps1 and GST-FgVps1^ΔPOC^ proteins were quantified using a Bradford Protein Assay Kit (Cat. P0006C, Beyotime, Shanghai, China) and then the GTPase activity was measured using a GTPase assay kit. GST was used for the negative control. Data were collected from three biological repeats.

## Author contributions

Conceptualization, ZHW and WHZ; methodology, XC, XMC and WHZ; investigation, XC, XMC, YFL, XT and ZYF; visualization, XC, ZHW and WHZ; Writing – Original Draft, XC; Writing – Review & Editing, XC, XMC, YSA, HWZ, ZHW and WHZ; resources and funding acquisition, XMC and WHZ. XC and XMC contributed equally to this work.

## Acknowledgements

This research was supported by the National Natural Science Foundation of China (32272481, 32122071), the Postdoctoral Fellowship Program of CPSF (GZC20230448). The authors declared no financial or other potential conflict of interest.

