## Supplemental figure S1-S9, Supplemental table S1-S3 for "Unraveling fungal endolysosomal network as a potential target for effective disease control"

### Supporting information

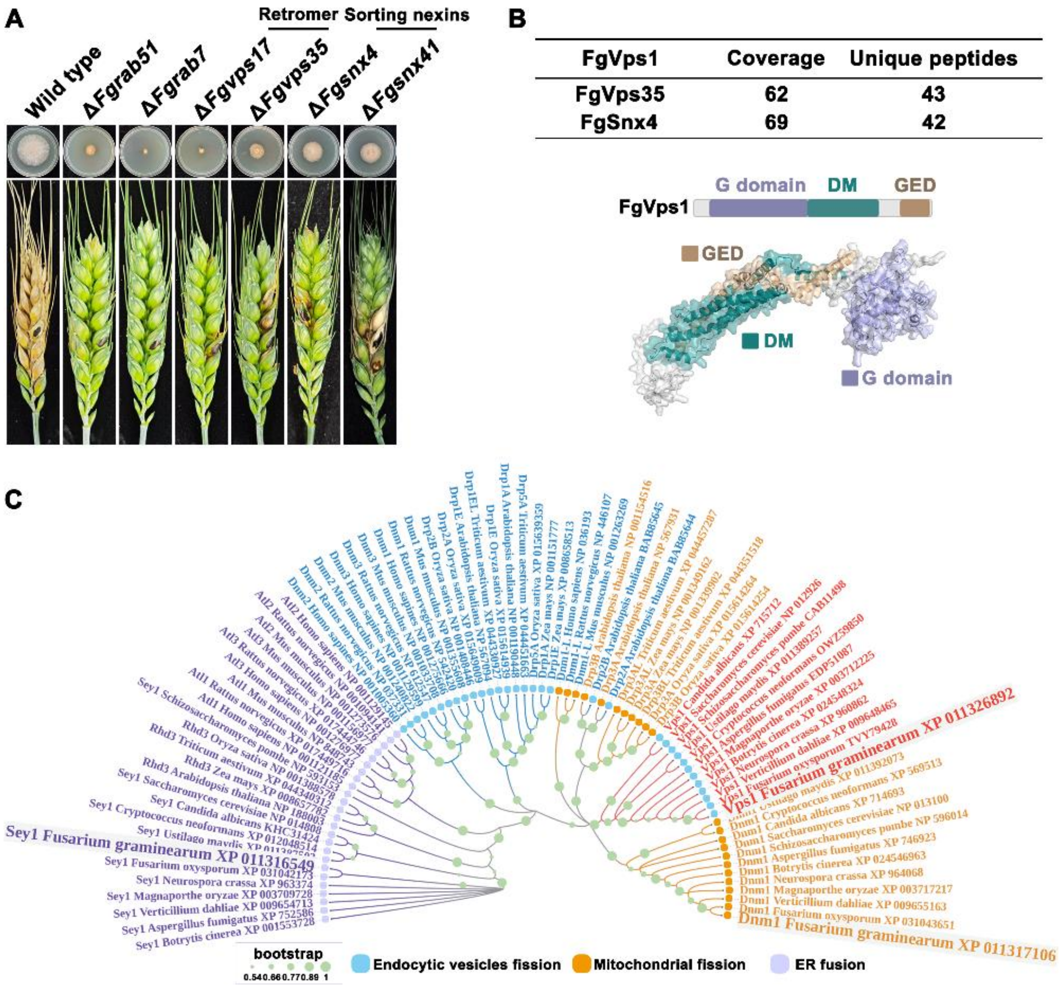


**Fig. S1 Identification of the dynamin-like GTPase FgVps1**

A. Growth and pathogenicity assays of the mutants of endolysosomal network components. B. FgVps1 is highly enriched in the mass spectrometry data of FgVps35 and FgSnx4. C. Phylogenetic analysis of the dynamin family proteins.

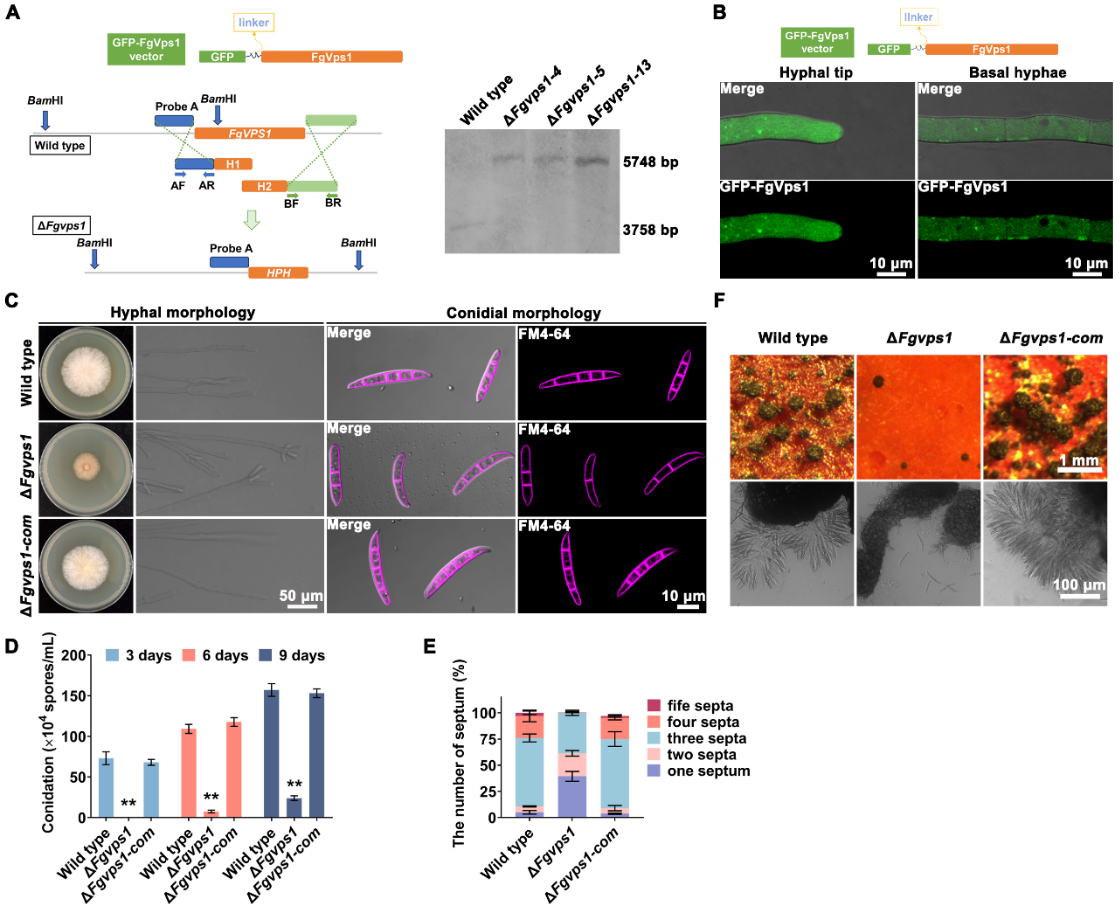


**Fig. S2 Phenotypic analysis of Δ*Fgvps1* mutant and its complemented strain**

A. Southern blot assay verifying the deletion of *FgVPS1* gene. B. Generation of the Δ*Fgvps1*-com complementation strain. For the construction of GFP-FgVps1 vector, a linker sequence (cgctccatcgccacgatg) was insert into the GFP fragment and *FgVPS1* ORF region to ensure the proper folding of FgVps1. The resulting vector was transformed into the Δ*Fgvps1* protoplast and then the transformants were screen under confocal microscopy. The GFP-FgVps1 signal was detected in the cytosol, as punctate structures at the hyphal tip and bubble-like structures in basal hyphae. C. Vegetative growth and conidial morphology assays of the indicated strains. D. Conidiation of the wild type, Δ*Fgvps1* and Δ*Fgvps1-com* strains at different time points (**$P$<0.01). E. Statistical analysis of the number of septa in wild type, Δ*Fgvps1* and Δ*Fgvps1-com* conidia. F. Analysis of sexual reproduction potential of the indicated strains.

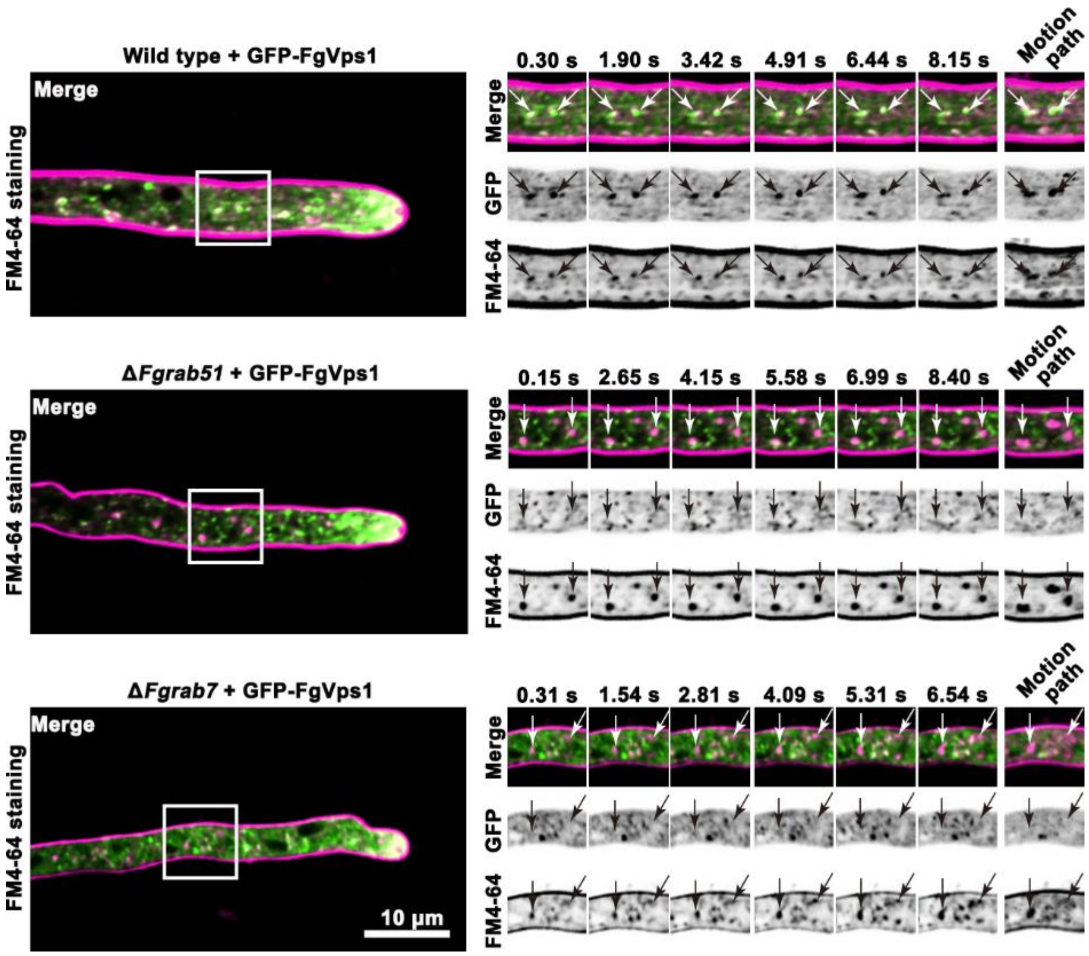


**Fig. S3 Loss of the Rab GTPases FgRab51 and FgRab7 impairs the association of FgVps1 with FM4-64-labelled endosomes**

For time-lapse imaging, the strains expressing GFP-FgVps1 were cultured on CM agar plates for 2 days and then stained with 10 μg/mL FM4-64. White boxes indicate the regions shown in the right panel. Arrows indicate the dynamics of FgVps1 in relation to FM4-64-labeled endosomal structures. Unlike the close association observed in the wild type, some FgVps1 puncta appeared spatially separated from the FM4-64-labeled endosomes in the Rab GTPase mutants.

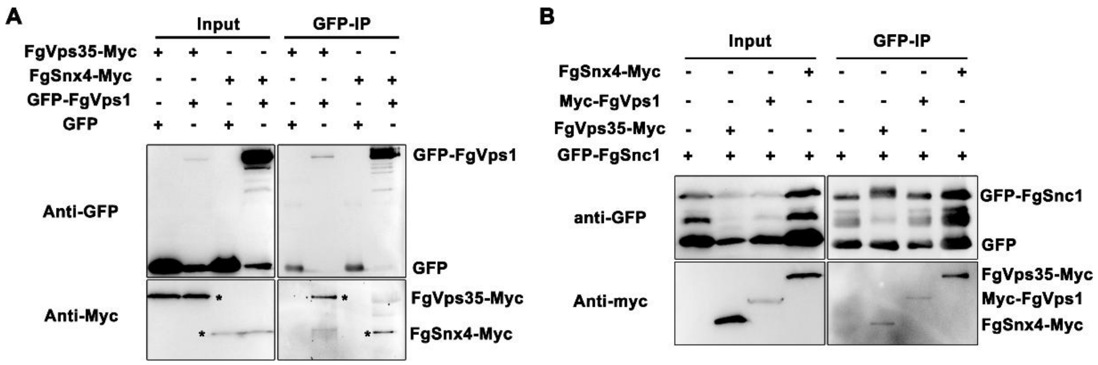


**Fig. S4 Co-immunoprecipitation (Co-IP) assays validating the interactions among FgVps1, FgSnx4, FgVps35 and FgSnc1**

A. *In vivo* validation of the interaction between FgVps1, FgSnx4 and FgVps35 through Co-IP assay.

B. Co-IP assay validating the interaction between FgVps1, FgSnx4 and FgVps35 with FgSnc1.

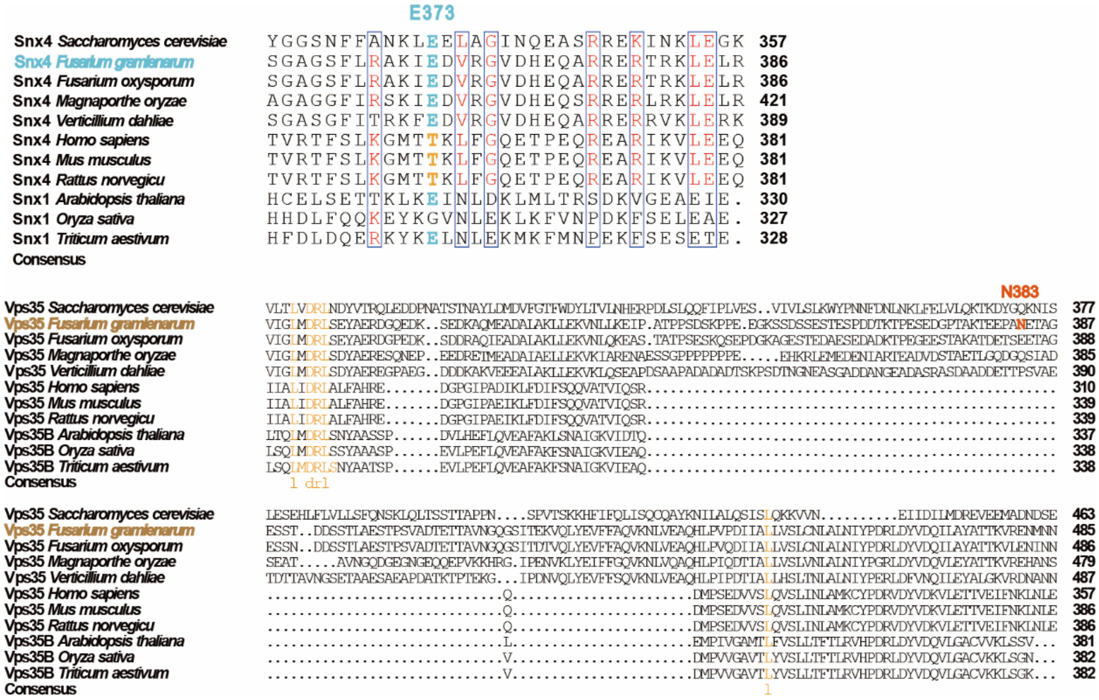


**Fig. S5 Sequence alignment of the FgSnx4 E373 and FgVps35 N383 sites across different species**

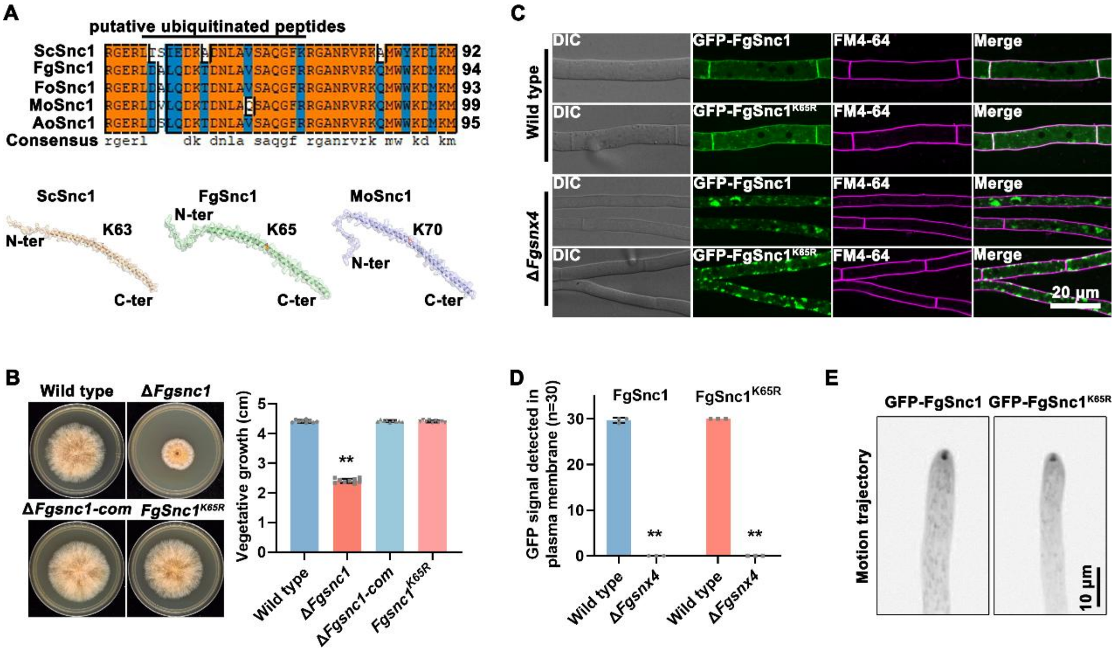


**Fig. S6 FgSnc1$^{K65}$ mutation does not affect its endosomal sorting**

A. Identification of FgSnc1 ubiquitination site by mass spectrometry. B. FgSnc1$^{K65R}$ mutation has no effect on the vegetative growth of *F. graminearum* (***P*<0.01). C. Localization of GFP-FgSnc1 and GFP-FgSnc1$^{K65R}$ in the basal hyphae of the wild type and Δ*Fgsnx4* mutant. D. Statistical analysis of FgSnc1 and FgSnc1$^{K65R}$ localization in the wild type and Δ*Fgsnx4* mutant (***P*<0.01). E. Motion trajectory of FgSnc1 and FgSnc1$^{K65R}$ in hyphal tips. A total of 57 images was taken at 500ms interval.

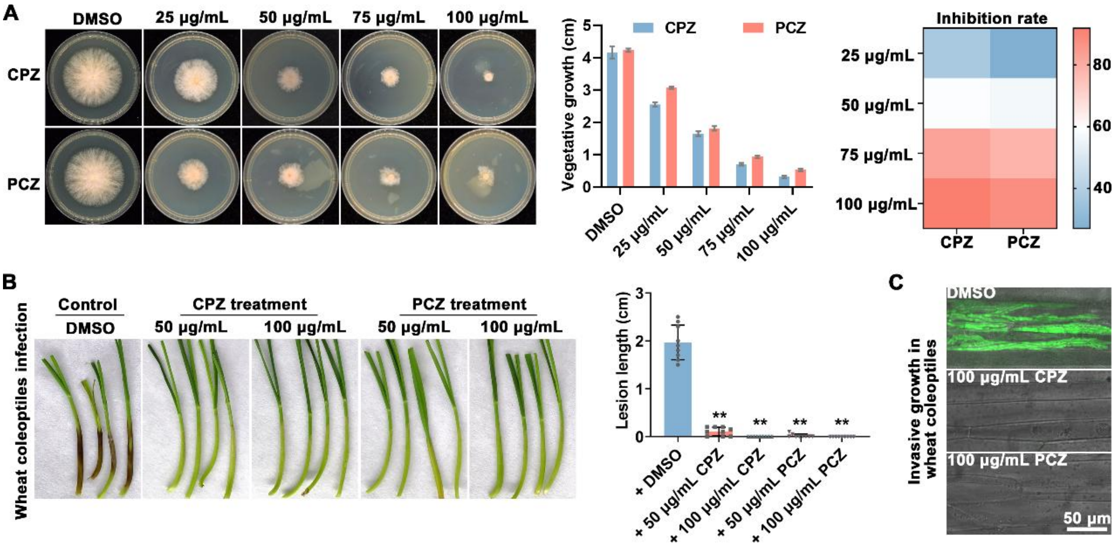


**Fig. S7 Treatment with phenothiazines affects the development and pathogenicity of *F. graminearum***

A. Evaluation of the effect of phenothiazines on fungal growth. Mycelial blocks of *F. graminearum* were inoculated on complete medium with or without phenothiazines for 2 days. B. Treatment with CPZ and PCZ impaired host invasion by *F. graminearum*. For wheat infection, 50 μL conidia suspensions ($10^6$ spores/mL from *F. graminearum* PH-1 strain was inoculated on wheat coleoptiles with or without CPZ or PCZ for 7 days. C. Investigation of *F. graminearum* invasive growth after treatment with phenothiazines. Conidia suspension from *F. graminearum* PH-1 strain expressing GFP was injected into wheat coleoptiles with CPZ or PCZ. DMSO treatment was used as control. No obvious invasive growth was detected at 3 dpi under treatment with phenothiazines.

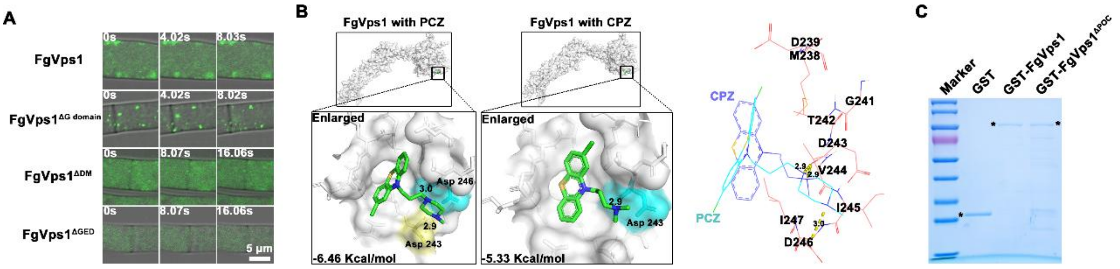


**Fig. S8 Identification of putative phenothiazine binding pockets in FgVps1**

A. Confocal examination for the effects of domain truncation on FgVps1 distribution. D. Molecular docking of prochlorperazine and chlorpromazine with FgVps1. The molecular docking of FgVps1with the phenothiazines was analyzed using AutoDock and further visualized by PyMOL. FgVps1 is depicted in surface representation (gray) while PCZ and CPZ are depicted in sticks representation. In the right panel, the binding pocket is depicted in lines (salmon), PCZ is represented in blue and CPZ is represented in light blue. C. Purification of GST, GST-FgVps1and GST-FgVps1$^{\Delta POC}$ recombinant proteins.

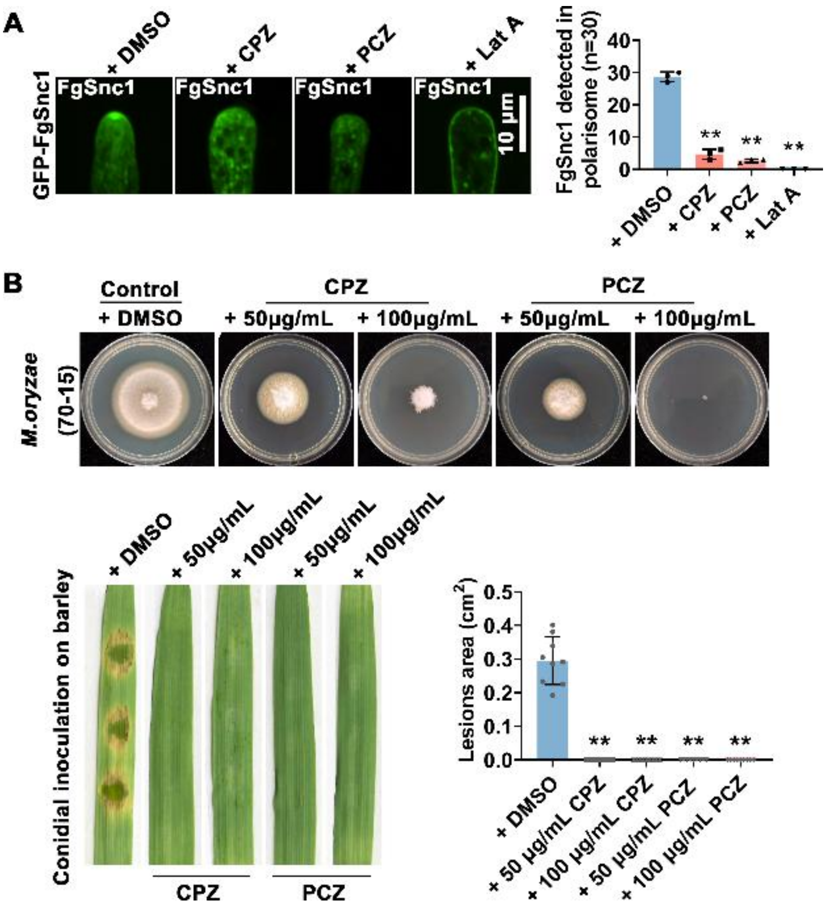


**Fig. S9 Disruption of actin polymerization affects the distribution of FgSnc1**

A. Effects of the actin inhibitor Latrunulin A (Lat A) on FgSnc1 distribution (***P*<0.01). To examine the impact of Lat A on FgSnc1 localization, hyphae were treated with 5 μg/mL Lat A for 30 minutes. B. Effects of PCZ and CPZ treatment on *M. oryzae* growth and host infection. For vegetative growth, the *M. oryzae* wild type strain 70-15 was inoculated on CMII medium for 7 days. For the pathogenicity on of *M. oryzae* on barley leaves, 10 μL conidia suspension was inoculated on barley leaves with PCZ or CPZ treatment for 5 days (***P*<0.01).

**Table S1 Strains used in this study**

| Strains | Description | References |
|---|---|---|
| PH-1 | Wild type strain of the *Fusarium graminearum* | [1] |
| Δ*Fgvps1* | *FgVPS1* deletion mutant at PH-1 background | This study |
| FgVps1$^{\Delta POC}$ | Δ*Fgvps1* mutant expressing mCherry-FgVps1$^{\Delta POC}$ vector | This study |
| Δ*Fgvps1*-com | Δ*Fgvps1* mutant expressing GFP-FgVps1 vector | This study |
| FgVps1$^{\Delta G\ domain}$ | Δ*Fgvps1* mutant expressing GFP-FgVps1$^{\Delta G\ domain}$ vector | This study |
| FgVps1$^{\Delta DM}$ | Δ*Fgvps1* mutant expressing GFP-FgVps1$^{\Delta DM}$ vector | This study |
| FgVps1$^{\Delta GED}$ | Δ*Fgvps1* mutant expressing GFP-FgVps1$^{\Delta GED}$ vector | This study |
| Δ*Fgvps35* | *FgVPS35* deletion mutant at PH-1 background | [2] |
| Δ*Fgvps17* | *FgVPS17* deletion mutant at PH-1 background | [2] |
| Δ*Fgsnx4* | *FgSNX4* deletion mutant at PH-1 background | [3] |
| Δ*Fgsnx41* | *FgSNX41* deletion mutant at PH-1 background | [3] |
| FgSnx4$^{E373A}$ | Δ*Fgsnx4* mutant expressing FgSnx4$^{E373A}$-mScarlet vector | This study |
| FgVps35$^{N383A}$ | Δ*Fgvps35* mutant expressing FgVps35$^{N383A}$-mScarlet vector | This study |
| Δ*Fgsnc1* | *FgSNC1* deletion mutant at PH-1 background | [3] |
| FgSnc1$^{K65R}$ | Δ*Fgsnc1* mutant expressing GFP-FgSnc1$^{K65R}$ vector | This study |
| Δ*Fgrab51* | *FgRAB51* deletion mutant at PH-1 background | [4] |
| Δ*Fgrab7* | *FgRAB7* deletion mutant at PH-1 background | [4] |
| Δ*Fgvps36* | *FgVPS36* deletion mutant at PH-1 background | [5] |
| FgVps35-GFP+ mCherry-FgVps1 | Wild type PH-1 strain expressing FgVps35-GFP and mCherry-FgVps1 vectors | This study |
| FgSnx4-GFP+ mCherry-FgVps1 | Wild type PH-1 strain expressing FgSnx4-GFP and mCherry-FgVps1 vectors | This study |
| Δ*Fgvps1* FgVps35-GFP+ FgSnx4-mCherry | Δ*Fgvps1* mutant expressing FgVPS35-GFP and FgSnx4-mCherry vectors | This study |
| FgVps35-Myc+ FgSnx4-GFP | Wild type PH-1 strain expressing FgVps35-Myc and FgSnx4-GFP vectors | This study |
| GFP-FgRab51+ mCherry-FgVps1 | Wild type PH-1 strain expressing GFP-FgRab51 and mCherry-FgVps1 vectors | This study |
| GFP-FgRab7+ mCherry-FgVps1 | Wild type PH-1 strain expressing GFP-FgRab7 and mCherry-FgVps1 vectors | This study |
| Myc-FgVps1+ GFP-FgRab51 | Wild type PH-1 strain expressing Myc-FgVps1 and GFP-FgRab51 vectors | This study |
| Myc-FgVps1+ GFP-FgRab7 | Wild type PH-1 strain expressing Myc-FgVps1 and GFP-FgRab7 vectors | This study |
| Δ*Fgrab51+* GFP-FgVps1 | Δ*Fgrab51* mutant expressing GFP-FgVps1 vector | This study |
| Δ*Fgrab7+* GFP-FgVps1 | Δ*Fgrab7* mutant expressing GFP-FgVps1 vector | This study |
| Δ*Fgvps36+* GFP-FgVps1 | Δ*Fgvps36* mutant expressing GFP-FgVps1vector | This study |
| GFP-FgSnc1+ | Wild type PH-1 strain expressing GFP-FgSnc1 and | This study |

| | | |
|---|---|---|
| mCherry-FgCfem1 | mCherry-FgCfem1 vectors | |
| Δ*Fgvps1*+ mCherry-FgCfem1 | Δ*Fgvps1* mutant expressing mCherry-FgCfem1 vector | This study |
| Δ*Fgsnx4*+ mCherry-FgCfem1 | Δ*Fgsnx4* mutant expressing mCherry-FgCfem1 vector | This study |
| Δ*Fgsnc1*+ mCherry-FgCfem1 | Δ*Fgsnc1* mutant expressing mCherry-FgCfem1 vector | This study |
| GFP-FgSnc1 | ΔFgsnc1 mutant expressing GFP-FgSnc1 vector | [3] |
| Δ*Fgsnx4*+ GFP- FgSnc1 | Δ*Fgsnx4* mutant expressing GFP-FgSnc1 vector | [3] |
| Δ*Fgvps35*+ GFP- FgSnc1 | Δ*Fgvps35* mutant expressing GFP-FgSnc1 vector | This study |
| Δ*Fgvps1*+ GFP- FgSnc1 | Δ*Fgvps1* mutant expressing GFP-FgSnc1 vector | This study |
| FgVps1$^{\Delta POC}$ + GFP- FgSnc1 | FgVps1$^{\Delta POC}$ strain expressing GFP-FgSnc1 vector | This study |
| Δ*Fgvps36*+ GFP-FgSnc1 | Δ*Fgvps36* mutant expressing GFP-FgSnc1vector | This study |
| GFP-FgSnc1$^{K65R}$ | Δ*Fgsnc1* mutant expressing GFP-FgSnc1$^{K65R}$ vector | This study |
| Δ*Fgsnx4*+ GFP- FgSnc1$^{K65R}$ | Δ*Fgsnx4* mutant expressing GFP-FgSnc1$^{K65R}$ vector | This study |
| FgSnx4-Myc+ GFP-FgSnc1 | Wild type PH-1 strain expressing FgSnx4-Myc and GFP-FgSnc1 vectors | This study |
| FgVps35-Myc+ GFP-FgSnc1 | Wild type PH-1 strain expressing FgVps35-Myc and GFP-FgSnc1 vectors | This study |
| Myc-FgVps1+ GFP-FgSnc1 | Wild type PH-1 strain expressing Myc-FgVps1 and GFP-FgSnc1 vectors | This study |
| FgSnx4-GFP+ mCherry-FgVps36 | Wild type PH-1 strain expressing FgSnx4-GFP and mCherry-FgVps36 vectors | This study |
| FgVps35-GFP+ mCherry-FgVps36 | Wild type PH-1 strain expressing FgVps35-GFP and mCherry-FgVps36 vectors | This study |
| FgSnx4-GFP | Δ*Fgsnx4* mutant expressing FgSnx4-GFP vector | This study |
| FgVps35-GFP | Δ*Fgvps35* mutant expressing FgVps35-GFP vector | This study |
| Δ*Fgvps36*+ FgSnx4-GFP | Δ*Fgvps36* mutant expressing FgSnx4-GFP vector | This study |
| Δ*Fgvps36*+ FgVps35-GFP | Δ*Fgvps36* mutant expressing FgVps35-GFP vector | This study |

**Table S2 Primers used in this study**

| Primers | Sequences (5'to 3') | Application |
|---|---|---|
| FgVps1 AF | TCCCTTGTGACTACGAATG | Generation of Δ*Fgvps1* deletion mutant |
| FgVps1 AR | ttgacctccactagctccagccaagccTGATGGTTGCCACTGATG | |
| FgVps1 BF | gaatagagtagatgccgaccgcgggttACGAGGCTTCTTATCATCACG | |
| FgVps1 BR | CGCCGATAGGGAACAGAG | |
| FgVps1 OF | ATCGCCGTCGTAGGAAGT | |
| FgVps1 OR | TTGATGTCTGGCAGGGTCT | |
| FgVps1 UA | TAAATCTGGTAAATGGGCTTCC | |
| H853 | GACAGACGTCGCGGTGAGTT | |
| FgVps1 proF | agggaacaaaagctgggtaccAATTTCTCTTAGATAGTGTA | Construction of GFP-FgVps1 and mCherry-FgVps1 vectors |
| FgVps1 proR | tcctcgcccttgctcaccatCTTCAAGTATTATTTCCCAG | |
| GFPF | ATGGTGAGCAAGGGCGAGGA | |
| GFPR | CTTGTACAGCTCGTCCATGC | |
| FgVps1 OGF | gcatggacgagctgtacaagcgctccatcgccacgatgATGAGCGGCTCCCTTGTTGC | |
| FgVps1 OGR | tcagtaacgttaagtggatccTAATGGGTGACTGATTATAT | |
| Myc-FgVps1 Pro F | agggaacaaaagctgggtaccAATTTCTCTTAGATAGTGTA | Construction of Myc-FgVps1 vector |
| Myc-FgVps1 Pro R | gagatcagcttctgctccatcatcgtggcgatggagcgCTTCAAGTATTATTTCCCAG | |
| Myc-FgVps1 ORF | ATGGAGCAGAAGCTGATCTCAGAGGAGGACCTGAGCGGCTCCCTTGTTGCACA | |
| Myc-FgVps1 ORR | gccgaattcgatatcaagcttTAATGGGTGACTGATTATAT | |
| Vps1 GED R1 | gttcatgaacgacatcactgGTTCTCGCGCTCTGATAGAG | Construction of GFP-FgVps1$^{\Delta GED}$, GFP-FgVps1$^{\Delta DM}$, GFP-FgVps1$^{\Delta G\ domain}$, GFP-FgVps1$^{\Delta POC}$ and mCherry-FgVps1$^{\Delta POC}$ vectors |
| Vps1 GED F2 | ctctatcagagcgcgagaacCAGTGATGTCGTTCATGAAC | |
| Vps1 DMR1 | ttgtaccgctcgttcaccatGTCAGTTCCCTCGTCCATCA | |
| Vps1 DMF2 | tgatggacgagggaactgacATGGTGAACGAGCGGTACAA | |
| Vps1 G R1 | AGGGTTGTTGACACCGACAG | |
| Vps1 G F2 | ctgtcggtgtcaacaaccctGACATCAAGGCCCGCATCAG | |
| Poc2 R1 | ggaatgactcggttggacaaCAGATCAACCTTGGTCAGGA | |
| Poc2 F2 | tcctgaccaaggttgatctgTTGTCCAACCGAGTCATTCC | |

| | | |
|---|---|---|
| FgVps1 PULLF | gatctggttccgcgtggatccATGAGCGGCTCCCTTGTTGC | Construction of GST-FgVps1 |
| FgVps1 PULLR | ctcgagtcgacccgggaattcTCACTGCACTTGGCTGACAA | |
| FgVps35 MycF | agggaacaaaagctgggtaccTTGCGGATCAGTCTGAGGTA | Construction of FgVps35-Myc vector |
| FgVps35 MycR | gccgaattcgatatcaagcttTCACAGGTCCTCCTCTGAGATCAGCTTCTGCTCGTTCGGTGTCAGCACAATAC | |
| FgVps35 GF | agggaacaaaagctgggtaccTTGCGGATCAGTCTGAGGTA | Construction of FgVps35-GFP and FgVps35$^{N383A}$-mScarlet vector |
| FgVps35 GR | gcccttgctcaccataagcttGTTCGGTGTCAGCACAATAC | |
| N383A R1 | CTACTTTCACCGGCAGTTTCGGCGGCAGGCTCCTCCGTTTTGG | |
| N383A F2 | CCAAAACGGAGGAGCCTGCCGCCGAAACTGCCGGTGAAAGTAG | |
| E373A R1 | GTGCGCTCGCGACGGGCTTGCGCGTGGTCTACGCCGCGGACAT | Construction of FgSnx4-GFP and FgSnx4$^{E373A}$-mScarlet vector |
| E373A F2 | ATGTCCGCGGCGTAGACCACGCGCAAGCCCGTCGCGAGCGCAC | |
| FgSnx4 GF | agggaacaaaagctgggtaccCCGACATGATGCTCCGATAC | |
| FgSnx4 GR | gcccttgctcaccataagcttAGCACTCGTGATACCCTGCT | |
| FgSnx4 ADF | gacgtaccagattacgctcatATGACGGCAACCGAACAGCA | Construction of the FgSnx4 yeast two hybrid vectors |
| FgSnx4 ADR | tatcgatgcccacccgggtggaaCTAAGCACTCGTGATACCCT | |
| FgSnx4 BDF | ctgatctcagaggaggacctgcatATGACGGCAACCGAACAGCAA | |
| FgSnx4 BDR | cgctgcaggtcgacggatccccgggaaCTAAGCACTCGTGATACCCT | |
| FgVps35 ADF | gacgtaccagattacgctcatATGGCCACTCCAGCTCCCCC | Construction of the FgVps35 yeast two hybrid vectors |
| FgVps35 ADR | tatcgatgcccacccgggtggaaTCAGTTCGGTGTCAGCACAA | |
| FgVps35 BDF | ctgatctcagaggaggacctgcatATGGCCACTCCAGCTCCCCC | |
| FgVps35 BDR | cgctgcaggtcgacggatccccgggaaTCAGTTCGGTGTCAGCACAA | |
| FgSnc1 ProF | agggaacaaaagctgggtaccTTCTATAGGGTCTTATATAG | Construction of GFP-FgSnc1$^{K65R}$ vectors |
| FgSnc1 ProR | tcctcgcccttgctcaccatTTTGGCTGTGTTTTGTGAAG | |
| FgSnc1 OGF | gcatggacgagctgtacaagATGCCTGAGCAGGAAGCCCC | |
| FgSnc1 OGR | tcagtaacgttaagtggatccCGTTCTTCGCCAAAGTATAT | |

| | | |
|---|---|---|
| FgSnc1 K65R R1 | CCTATCCTGCAAAGCGTCGA | |
| FgSnc1 K65R F2 | tcgacgctttgcaggataggACCGACAACTTGGCTGTTTC | |
| FgVps36 pro F | agggaacaaaagctgggtaccCATTACAGGCAAAATCGGTA | Construction of the mCherry-FgVps36 vector |
| FgVps36 pro R | tcctcgcccttgctcaccatCCGTCATGGAGGTCAAAGTA | |
| FgVps36 ORF | gcatggacgagctgtacaagATGTTTCTGAAACACATCGA | |
| FgVps36 ORR | tcagtaacgttaagtggatccCTTGCTACCTACTTATTCAT | |
| FgHse1 ADF | gacgtaccagattacgctcatATGTTCCGAGCCTCAGCAGC | Construction of the yeast two hybrid vectors of the ESCRT components |
| FgHse1 ADR | tatcgatgcccacccgggtggaaCTAATAAACCCCGCCACTTT | |
| FgVps27 ADF | gacgtaccagattacgctcatATGATGAGTTGGTGGTCGTC | |
| FgVps27 ADR | tatcgatgcccacccgggtggaaTTAAAACTCAATCAGGGCCT | |
| FgVps23 ADF | gacgtaccagattacgctcatATGCCCGTCCAGCAGCATGT | |
| FgVps23 ADR | tatcgatgcccacccgggtggaaTCACGCAGCCAGGCCCATTC | |
| FgVps28 ADF | GACGTACCAGATTACGCTCATATGATCCCACGACAAGGCTA | |
| FgVps28 ADR | tatcgatgcccacccgggtggaaTCAGGTCAAAGTCCTTCTAA | |
| FgVps22 ADF: | gacgtaccagattacgctcatATGTCTCGTAAAGGCGTAGG | Construction of the yeast two hybrid vectors of the ESCRT components |
| FgVps22 ADR | tatcgatgcccacccgggtggaaTCAACCTCCCTCCTCAGGAT | |
| FgVps25 ADF | gacgtaccagattacgctcatATGCTACAGAGCCTAGAACC | |
| FgVps25 ADR | tatcgatgcccacccgggtggaaTTAGAAGAATTTCACGCCTA | |
| FgVps36 ADF | gacgtaccagattacgctcatATGTTTCTGAAACACATCGA | |
| FgVps36 ADR | tatcgatgcccacccgggtggaaCTATAAGAGGCCGCTTGCTT | |
| FgVps20 ADF | gacgtaccagattacgctcatATGGGTGGGAATGCGAGTAA | |
| FgVps20 ADR | tatcgatgcccacccgggtggaaTCATGCTGCAAGCATAGCTG | |

| | |
|---|---|
| FgSNF7 ADF: | gacgtaccagattacgctcatATGTGGGGTTGGTTCGGTGG |
| FgSNF7 ADR | tatcgatgcccacccgggtggaaTCACATAGCCATCTCGGCCT |
| FgVps24 ADF | gacgtaccagattacgctcatATGGAAACGTTTAAATCTCT |
| FgVps24 ADR | tatcgatgcccacccgggtggaaCTAACTGCGAAGCGCCTCTA |
| FgVps2 ADF | gacgtaccagattacgctcatATGAATATCCTGGAATGGGC |
| FgVps2 ADR | tatcgatgcccacccgggtggaaCTATTTCCGCAGACTGTCCA |

**Table S3 Putative FgVps1 interacting proteins identified by mass spectrometry**

| Accession | Coverage (%) | | |
|---|---|---|---|
| | **Repeat 1** | **Repeat 2** | **Repeat 3** |
| FgVps1 (FGSG_07172) | 88 | 90 | 93 |
| FgRab51 (FGSG_05501) | 7 | 43 | 34 |
| FgRab7 (FGSG_05141) | 20 | 49 | 49 |
| FgSnx4 (FGSG_09157) | 32 | 15 | 11 |
| FgSnx41 (FGSG_06950) | 31 | 20 | 21 |
| FgVps35 (FGSG_02756) | 20 | 7 | 4 |
| FgVps26 (FGSG_01155) | 16 | 25 | 19 |
| FgVps29 (FGSG_01552) | 34 | 18 | - |
| FgVps5 (FGSG_02011) | 33 | 2 | 27 |
| FgVps17 (FGSG_09600) | 3 | 21 | 23 |
| FgSnc1 (FGSG_08537) | 14 | 33 | 31 |
